# Dissociating encoding of memory and salience by manipulating long-term synaptic potentiation

**DOI:** 10.1101/2022.01.04.474865

**Authors:** Konstantin Kaganovsky, Mark H. Plitt, Renzhi Yang, Richard Sando, Lisa M. Giocomo, Jun B. Ding, Thomas C. Südhof

## Abstract

Neural codes are thought to be reorganized during memory formation by long-term potentiation (LTP) of synapses. Here, using a novel approach for selectively blocking LTP, we found that eliminating LTP in hippocampal or striatal circuits only produces limited effects on learning and memory. To reconcile the discrepancy between the large physiological effect of blocking LTP and the absent effect on learning, we studied how LTP impacts neuronal computations in the hippocampus using *in-vivo* Ca^2+^-imaging. Contrary to current conceptual frameworks, we found that hippocampal CA1-region LTP is not required for accurate representations of space in hippocampal neurons, but rather endows these neurons with reward- and novelty-coding properties. Thus, instead of driving formation of cognitive maps and memory engrams, CA1-region LTP incorporates salience information into cognitive representations.

**One-Sentence Summary:** A novel approach for studying long-term potentiation reveals its surprising and selective role in salience encoding

## Main Text

Learning is a fundamental process that enables an animal to adapt its behavior to changing environments. The discovery of long-term potentiation (LTP) in the hippocampus provided the first evidence for a physiological mechanism that could update neural activity patterns to support learning (*1, 2*). Since then, decades of work have demonstrated that proteins required for LTP are also essential for normal neural representations (*3*) and learning (*4*). Most notably, NMDA-receptors (NMDARs) are required for hippocampal LTP, accurate spatial coding, and learning (*5–7*). These results reasonably suggested that LTP is part of the learning mechanism. While this conclusion was formed based on the best available approaches at the time, it is now known that NMDARs are involved in multitudinous synaptic functions, including basic synaptic transmission at most excitatory synapses. Therefore, the original manipulations that disrupted both LTP and learning also impaired other synaptic processes, suggesting that the behavioral effects may have been caused by impairments in neuronal processes independent of LTP (*8–11*).

One promising approach for abolishing LTP while preserving basal synaptic properties is to prevent the activity-dependent insertion of AMPA-receptors (AMPARs) (*12–14*). Increasing evidence implicates a postsynaptic SNARE complex in AMPAR insertion during LTP (*15–20*). We previously showed that shRNA-mediated knockdown of a critical component of this postsynaptic SNARE complex, *Syntaxin-3* (*Stx3*), in CA1-region neurons impaired LTP while sparing basal synaptic transmission (*16*, *17,* but see (*21*) for an opposing view). In the current set of experiments, we explored whether we could use *Stx3* as a molecular handle to selectively probe the role of LTP in population coding and memory formation.

To validate the selective role of *Stx3* in LTP, we stereotactically injected AAVs encoding inactive (ΔCre) or active Cre-recombinase (Cre; both tagged with EGFP) into the CA1-region of the dorsal hippocampus of *Stx3^fl/fl^* mice, and performed whole-cell patch-clamp recordings from EGFP+ neurons (Fig1A, *22*). Postsynaptic deletion of *Stx3* in the CA1-region abolished LTP induced by tetanic stimulation of CA3-region axons (Fig 1B-C), but had no effect on basal synaptic transmission, the AMPAR/NMDAR ratio, presynaptic release probability, gross dendritic morphology, or spine density (Fig 1D, S1).

**Fig. 1.**
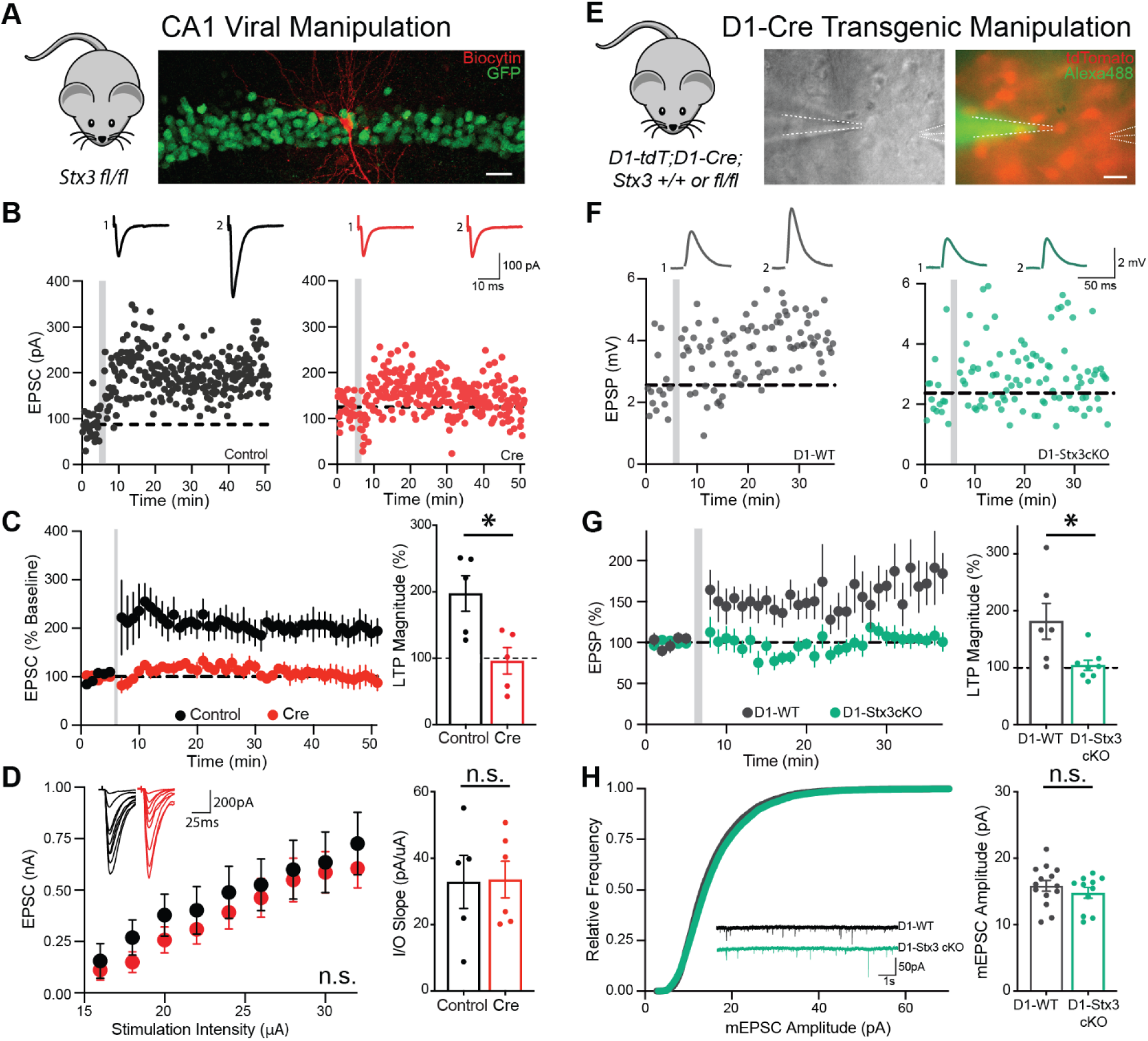
**A)** Schematic of electrophysiological experiments in the CA1 region. Confocal image shows EGFP-Cre fusion (green) and streptavidin Alexa555-biocytin staining after slice fixation (red). Scale bar: 50 µm **B)** Sample LTP experiment in CA1 pyramidal neurons from control and Cre cells. Gray bar indicates LTP pairing in this and subsequent panels. EPSCs during baseline (1) and last 5 min (2) are shown above the graph. **C)** *Stx3* deletion abolishes LTP. *Left-* summary plot of the normalized EPSC amplitude vs. time. *Right-* summary graph of the normalized EPSC amplitude 5 min before LTP induction and 41-45 min after LTP induction. [N=5 control cells, 5 Cre cells. Unpaired t-test: p=0.016] **D)** *Stx3* deletion does not impair basal synaptic strength as demonstrated by input/output measurements. *Left-* summary plot of absolute EPSC amplitudes as a function of the stimulus strength, with representative traces shown on top. *Right-* summary graph of the input/output slope. [N=5 control cells, 6 Cre cells. *Left-* rmANOVA: stimulation intensity main effect p<10^−5^, virus main effect p=0.54, interaction p=0.963; *Right-* unpaired t-test: p=0.943] **E)** Schematic of electrophysiological experiments in striatal neurons of adult mice. Images show DIC (left) and fluorescence views (right) of an acute slice. The patch pipette was filled with Alexa488 as one quality control measure of the perforated patch configuration (scale bar: 20 µm). **F)** Sample perforated-patch spike-timing-dependent LTP experiment. Only D1-tdT+ neurons were recorded from mice also carrying D1-Cre;*Stx3^+/+^* (left) or D1-Cre;*Stx3^fl/fl^* (right) alleles. EPSPs during baseline and last 5 minutes are shown as insets. **G)** *Stx3* deletion abolishes LTP in D1R^+^ neurons, showing that the role of *Stx3* in LTP is consistent across the induction protocols, cell-types, and brain regions tested. Summary graph of LTP as a function of time (left) and LTP magnitude (right). [N=6 D1-WT cells, 8 D1-*Stx3cKO* cells. Unpaired t-test: p=0.02] **H)** *Stx3* deletion does not affect basal synaptic strength in striatal D1R^+^ neurons as demonstrated by mEPSC amplitude. *Left-* cumulative frequency of mEPSC amplitude; inset: representative mESPC traces. *Right-* summary plot of mEPSC amplitude. [N=14 D1-WT cells, 11 D1-*Stx3cKO* cells. Unpaired t-test: p=0.365] * indicates p<0.05

The complete block of CA1-region LTP achieved by the genetic deletion of *Stx3* strengthens the previous shRNA knockdown results, but does *Stx3* have a broader role in LTP? To address this question, we studied whether *Stx3* is also required for LTP using a different induction procedure in a different cell-type and brain region. We deleted *Stx3* from dopamine D1-receptor-expressing cells by breeding *Stx3^fl/fl^* mice into a D1-cre^+/-^;D1-tdT^+/-^ background (D1-WT, D1-Stx3cKO, Fig S2A-C). We then studied spike-timing-dependent LTP in the striatum (Fig 1E) (*23*). Similar to CA1-region LTP, we found that *Stx3* is also required for spike-timing-dependent LTP in striatal D1R^+^ neurons (Fig 1F-G). Again, we confirmed that *Stx3* is not essential for basal synaptic transmission as measured by miniature excitatory postsynaptic currents (mEPSC) and miniature inhibitory postsynaptic currents (mIPSC) frequency and amplitude (Fig 1H, S2D-F). These data provide evidence for a general role of *Stx3* in LTP, regardless of the LTP induction protocol and brain region. Thus, the *Stx3^fl/fl^* mouse serves as a broadly applicable tool for studying the relationship between LTP, neural coding, and behavior.

Given the widely accepted idea that LTP is the cellular substrate of learning (*24*), we hypothesized that deletion of *Stx3* from CA1 or D1R^+^ cells would abolish learning mediated by hippocampal or striatal circuits. We deleted *Stx3* in dorsal CA1-region neurons by bilateral injections of AAV-Cre in *Stx3^fl/fl^* mice (“Cre mice”), with AAV-ΔCre injections as a control (“control mice”). We achieved nearly complete coverage of the dorsal CA1 region (Fig S3A-C). To our surprise, blocking LTP with the *Stx3* deletion did not impair any measure of learning or memory in cued or contextual fear-conditioning (Fig 2A, S3D-G), or in AMPAR subunit GluR1- (*25*) and NMDAR-dependent (*26*) trace fear-conditioning (Fig 2B,S3H-J). With encoding and retrieval intact, we next tested the role of CA1 LTP in long-term memory consolidation. We chose a time point in which the expression of long-term contextual fear memory is dependent on the dorsal hippocampus (*27*). However, 10 days after trace fear-conditioning, Cre and control mice behaved identically in the original conditioning context (Fig 2B). These data indicate that CA1 LTP is dissociable from the encoding, consolidation, and retrieval of fear memory across multiple procedures.

**Fig. 2.**
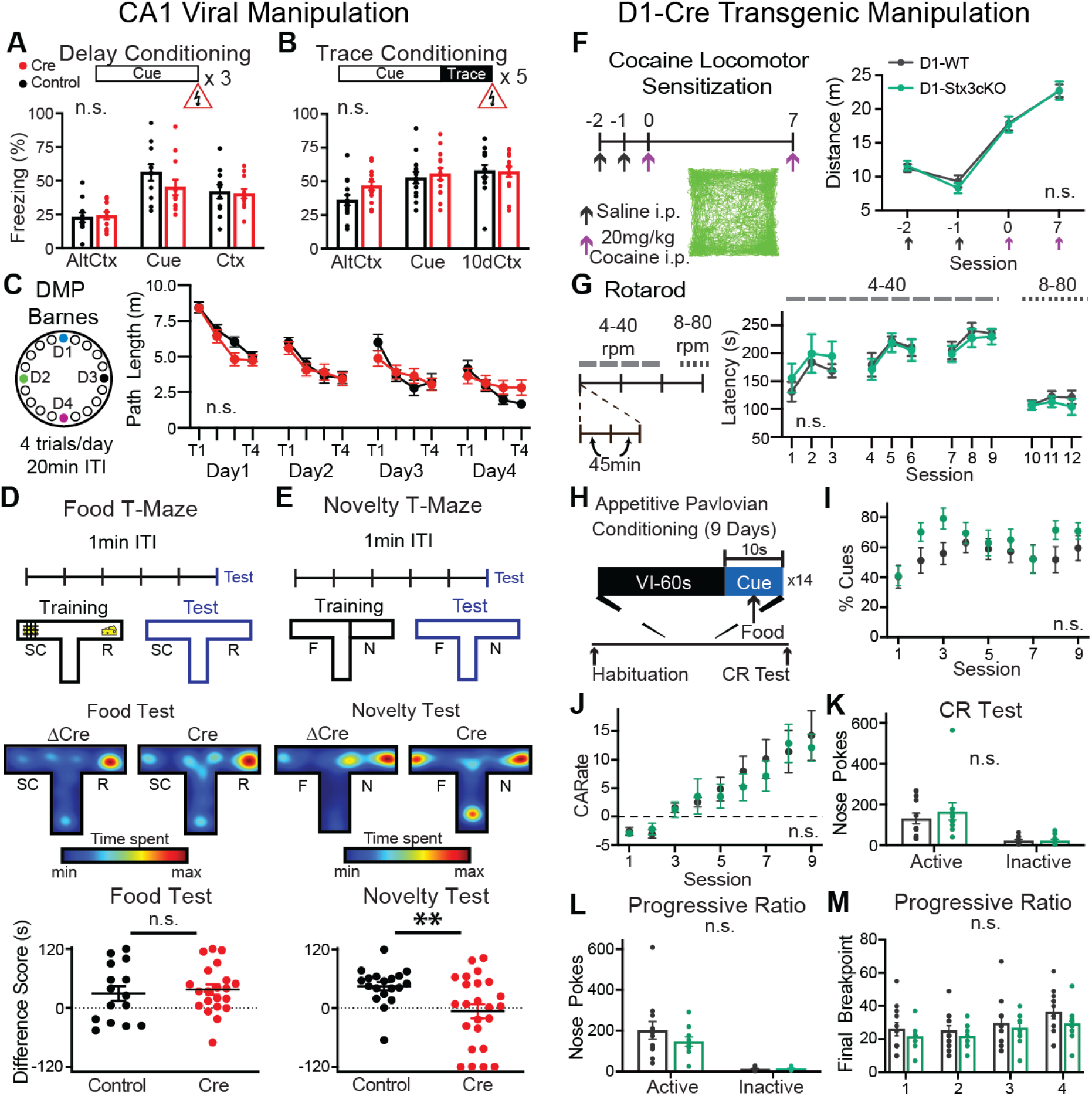
LTP is not required for common learning tasks known to depend on CA1-region/striatal neural activity and NMDA-receptor function. **A-B**) CA1 LTP is not required for fear conditioning across multiple procedures. *Top-*Schematic of the conditioning procedure. **A**) Summary plot of freezing during the altered context (AltCtx), cue, and the training context (Ctx). [N=11 control mice, 12 Cre mice. rmANOVA: test condition main effect p<10^−5^, virus main effect p=0.496, interaction p=0.147]. **B**) Summary plot of freezing during AltCtx, cue, and the training context 10 days after conditioning (10dCtx). [N=14 control mice, 14 Cre mice. rmANOVA: test condition main effect p<10^−5^, virus main effect p=0.375, interaction p=0.112] **C**) Delayed matching-to-place (DMP) Barnes Maze test reveals that CA1 LTP is dispensable for spatial learning and daily re-learning. *Left-* test design. *Right-* summary plot of average path length before entering the escape port. [N=19 control mice, 17 Cre mice. rmANOVA: trial main effect p<10^−5^, virus main effect p=0.741, interaction p=0.362] **D-E**) CA1 LTP is required for novelty-, but not reward-driven, spatial learning. *Top-* assay design. *Middle-* Representative mouse during test. *Bottom-* Dotted line indicates equal time spent in both arms. **D**) Food T-Maze difference score = (time in the rewarded arm)-(time in the scent control arm). [N=15 control mice, 21 Cre mice. Unpaired t-test: p=0.66]. **E**) Difference score = (time in N)-(time in F). [N=19 control mice, 24 Cre mice. Unpaired t-test: p=0.007] **F)** LTP in D1R^+^ neurons is not required for basal locomotion nor cocaine locomotor sensitization. *Left-* design of the experiment and representative track. *Right-* summary plot of the distance traveled over each 1hr session. [N=13 D1-WT mice, 15 D1-*Stx3cKO* mice. rmANOVA: session main effect p<10^−5^, genotype main effect p=0.844, interaction p=0.862] **G)** The accelerating rotarod assay demonstrates that LTP in D1R^+^ neurons is not involved in motor learning. *Left-* experimental design. *Right-* summary graph of the fall-off latencies. [N=6 D1-WT mice, 6 D1-*Stx3cKO* mice. mixed effects ANOVA: trial main effect p<10^−5^, genotype main effect p=0.997, interaction p=0.732] **H-J**) The Pavlovian conditioning task demonstrates that LTP in D1R^+^ neurons does not affect learning a cue-reward association. **H**) Experimental design. **I**) The % of trials in which a port entry (PE) occurred during the cue plotted by session. [N=12 D1-WT mice, 11 D1-*Stx3cKO* mice. rmANOVA: session main effect p=0.004, genotype main effect p=0.204, interaction p=0.293]. **J**) The conditioned approach rate [CArate = conditioned stimulus PE rate - ITI PE rate] is plotted as a function of session and demonstrates both groups learned to discriminate PE during the conditioned stimulus vs ITI. [N=12 D1-WT mice, 11 D1-*Stx3cKO* mice. rmANOVA: session main effect p<10^−5^, genotype main effect p=0.753, interaction p=0.933] **K**) The conditioned reinforcement test demonstrates that LTP in D1R^+^ neurons is not required for the same cue used in pavlovian conditioning to gain reinforcing properties in an operant setting. [N=12 D1-WT mice, 11 D1-*Stx3cKO* mice. rmANOVA on log-transformed data: port main effect p<10^−5^, genotype main effect p=0.629, interaction p=0.44] **L-M**) LTP in D1R^+^ neurons is not required to adaptively increase operant responding for a food pellet under a progressive ratio schedule of reinforcement. **L**) Nose-pokes into the active and inactive ports are shown, averaged over test sessions. [N=12 D1-WT mice, 10 D1-*Stx3cKO* mice. rmANOVA: port main effect p<10^−5^, genotype main effect p=0.321, interaction p=0.29]. **M**) The highest response requirement completed (i.e. breakpoint) across days is plotted. [N=12 D1-WT mice, 10 D1-*Stx3cKO* mice. rmANOVA session main effect p<10^−5^, genotype main effect p=0.322, interaction p=0.674] * indicates p<0.05 ** indicates p<0.01

It is possible that the intensity of the stimulus in fear-conditioning masked subtle changes in the animals’ ability to learn after LTP ablation. To overcome this, we tested spatial memory in a delayed matching-to-place version of the Barnes maze. We trained mice to escape through a particular hole for 4 trials/day with 20 minutes between trials, and changed the escape port’s quadrant daily (Fig 2C). Critically, both hippocampal lesions and hippocampal NMDAR antagonism dramatically impair learning in the delayed matching-to-place Morris Water Maze task with a 20 minute inter-trial-interval (*28*). Again, we found no effect of the *Stx3* deletion in the CA1-region on this spatial learning procedure: both groups showed identical significant decreases in path length (Fig 2C) and latency (Fig S4) over trials.

The delayed matching-to-place Barnes and fear-conditioning tasks both provided aversive stimuli to drive learning. We, therefore, opted for spatial learning tasks that rely on appetitive stimuli. We allowed food-restricted mice free access to a T-Maze, baited one arm with food, and baited the other arm with a scent-control (Fig 2D). Cre and control mice did not differ during training (Fig S5B) or during the test of arm preference in the Food T-Maze (Fig 2D, extinction conditions), indicating that CA1 LTP is also dispensable for forming a memory of a reward location.

Building upon pioneering work studying global GluR1 KO mice, which exhibit impairments in basal synaptic transmission, LTP, motor skills, and learning (*12, 29–31*), we next studied the role of CA1-region *Stx3* in a spatial novelty task. In this T-Maze, we allowed mice to explore 2 of 3 arms during training (Fig S5E-F), and the full T-Maze during the test (Fig 2E). During the test, control mice preferred the previously unseen arm of the T-Maze, while Cre-injected mice explored the maze at chance levels (Fig 2E). This directly confirms the hypothesis that CA1 LTP is necessary for short-term spatial novelty memory (*31*), and indicates that viral ablation of CA1 LTP is sufficient to affect learning and memory (i.e. these data serve as a positive control for our manipulation).

Taken together, these data demonstrate a dissociation between CA1 LTP and cued/contextual fear learning (with or without a trace period), contextual discrimination, and multiple forms of spatial learning. They also reveal the necessity of CA1 LTP for spatial novelty preference, hinting at a role for CA1 LTP in the neural code that drives this behavior.

While the hippocampus is a hub in the medial temporal lobe’s cognitive learning system, the basal ganglia make up a separate learning system that supports motor, stimulus-response (S-R), and reinforcement learning (*32, 33*). The striatum integrates excitatory inputs with instructive dopaminergic signals, an ideal architecture for Hebbian coincidence detection to exert a role on behavior (*34*). Indeed, cocaine exposure changes excitatory synaptic strength onto dopamine D1 receptor-expressing (D1R^+^) neurons (*35*), a major cell-type within the striatum that can drive reinforcement (*36*). This drug-induced plasticity is thought to be required for the enhanced locomotor response to subsequent doses of cocaine, i.e. “cocaine-locomotor sensitization” (*37–39*). We, therefore, tested whether ablating LTP in D1R^+^ neurons by deletion of *Stx3* would block cocaine-locomotor sensitization. Regardless of whether the mice had normal LTP in D1R^+^ neurons, both groups exhibited a sensitized locomotor response to cocaine given one week earlier (Fig 2F), indicating a dissociation between LTP in D1R^+^ neurons and cocaine-locomotor sensitization. To validate this finding, we opted for a simpler motor learning task that requires NMDARs in the striatum and D1-receptor signaling during early learning: the accelerating rotarod (*40, 41*). Again, both groups learned the motor skill at a similar rate (Fig 2G), indicating that LTP in D1R^+^ neurons is not required for driving enhanced locomotion and motor skill learning.

The striatum is also critical for reward-related behaviors, and cortico-striatal synapses undergo potentiation during reward learning (*34*). Therefore, we probed the necessity of LTP in D1R^+^ neurons in Pavlovian reward learning (Fig 2H). While there are conflicting reports on the role of NMDARs in D1R^+^ neurons in Pavlovian conditioning (*42, 43*), we found that blocking LTP in D1R^+^ neurons by deleting *Stx3* did not affect learning this task (Figs 2I-J, S7A). Additionally, we did not observe an effect of abolishing LTP in D1R^+^ neurons on learning measured by instrumental/operant conditioning – the conditioned stimulus alone drove reinforcement of a learned behavior (i.e. conditioned reinforcement test, Figs 2K, S7B). Lastly, we allowed fully fed mice to operantly self-administer food without cues, and did not identify a difference between genotypes in either a self-administration procedure only requiring one response per reward, or a procedure with a progressively increasing requirement for each reward (Figs 2L-M, S7C-G). These data imply that learning a cue-food reward association does not require LTP in D1R^+^ neurons, nor does the ability to use food to reinforce behavior. Combining these behavioral results across brain regions, we find that using the *Stx3* deletion to eliminate LTP yields the surprising result that only a limited set of behaviors previously shown to depend on the hippocampus or striatum require LTP in these regions.

To better understand why we observed such selective and puzzling behavioral deficits in the absence of LTP, we turned to *in vivo* two-photon Ca^2+^-imaging of hippocampal neurons during head-fixed navigation in a virtual reality maze. This method enabled us to analyze the role of LTP in broader neural computations by recording the activity of large populations of CA1-region neurons (Figs 3A, S8 & 9, <u>Movie S1</u>; 250-1,840 cells imaged simultaneously). The neural representations of memory have been extensively studied in CA1 neurons and their synaptic partners, allowing contextualization of our results. Most notably, all hippocampal subfields contain “place cells” (*44–46*), a functionally defined cell class that fires action potentials specifically when an animal is in one or several locations within an environment. Within an environment, place cells tile the space via their place fields. As an animal moves between environments, place cells will turn on, turn off, move their location of peak firing, or change their peak firing rate (*47, 48*). These phenomena are collectively referred to as “remapping” and result in independent representations of space in different spatial contexts. Given that place cells are required for spatial navigation and memory (*49*), we studied the effects of LTP on the properties of these “cognitive maps”.

**Fig. 3.**
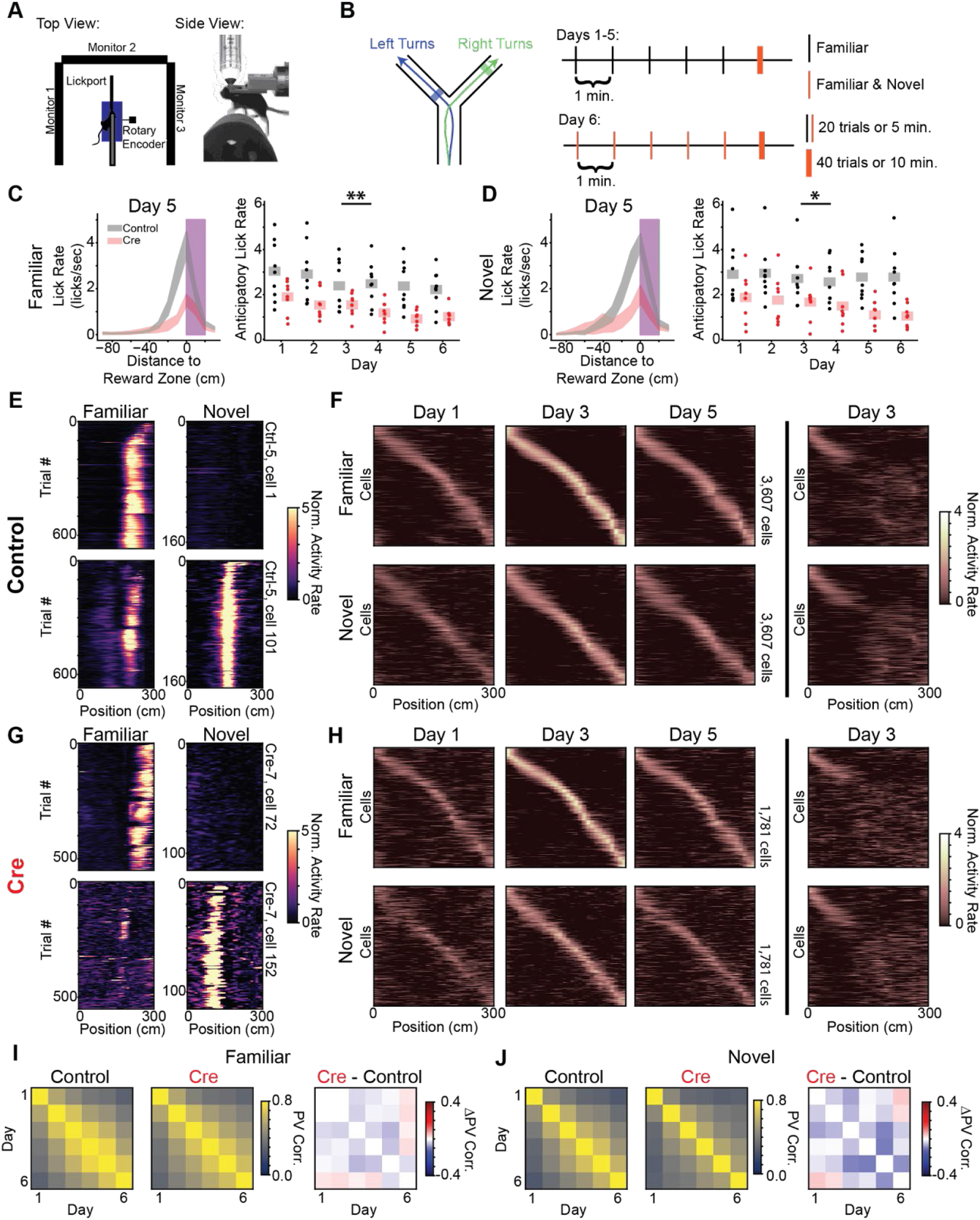
CA1-region LTP increases reward anticipation but is not required to form stable representations of position and context. **A)** Head-fixed virtual reality (VR) and two photon (2P) rig. **B)** Task design of virtual reality forced Y-Maze. *Left-*Track schematic. Arrows indicate the animals’ trajectory on left and right trials. Shaded regions indicate reward zones. *Right-*Training protocol. Days 1-5: 5 blocks of “familiar arm” trials (20 trials or 5 minutes, black, identity of familiar arm counterbalanced across animals), 1 block of randomly interleaved familiar and “novel arm” trials (40 trials or 10 minutes, orange). Day 6: familiar and novel arms randomly interleaved in all blocks. **C-D**) CA1 LTP is critical for predictive consummatory behavior. **C**) *Left -* peri-reward lick rate on familiar arm trials as a function of position on day 5 (red - Cre, black-control, magenta - reward zone). Data shown as across animal mean SEM. *Right -* Average familiar trial peri-reward lick rate on each day. Dots indicate the across trial average for each mouse. Shaded bars indicate across animal mean. This format is used for all summary graphs to follow. [N=9 control mice, 8 Cre mice, 6 days. Mixed effects ANOVA: virus main effect p=5.22×10^−4^, day main effect p=8.61×10^−6^, interaction p=0.55]. **D**) Same as (C) for novel arm trials. [N=9 control mice, 8 Cre mice, 6 days. Mixed effects ANOVA: virus main effect p=0.011, day main effect p=0.087, interaction p=0.263] **E-J**) LTP is dispensable for forming stable spatial representations and for remapping of place cells across arms of the maze. **E**) Example co--recorded stable place cells tracked over 6 days from a control mouse. Each subpanel shows trial x position activity rate of a single cell (rows) on familiar (left) and novel (right) trials. Place fields are stable over days and remap across trial types. **F**) Same phenomena observed in (E), but for all place cells tracked over all 6 recording days for familiar (top row) and novel (bottom row) trials. Z-scored trial-averaged activity rate is plotted for each cell on a subset of days. Cells are in the same order for each plot within a row and are sorted by the location of peak activity on odd-numbered trials from day 3. Day 3 heatmaps indicate average activity on even trials. All other heatmaps indicate the average of all trials. These heatmaps show a stable tiling of place cells across days. *Far right-*novel trial activity with cells sorted by familiar trial activity (top). Familiar trial activity with cells with cells sorted by novel trial activity (bottom). Place cells that code for the stem of the Y-Maze are visible at the beginning of the track. Place cells remap on the arms of the maze as indicated by the disorganized activity rate maps. **G-H**) Same as (E-F) for example Cre cells. Place cell representations from Cre mice are highly similar to those from Control mice. **I**) Across animal average day x day population vector correlation (PV corr.) on familiar trials for control (left) and Cre animals (middle) and the difference between groups (right, [N=9 control mice, 7 Cre mice. pairwise t-tests Holm corrected p>0.05]). Each animal’s population vector is calculated using the across trial average spatial activity rate map for each cell on each day. **J**) Same as (I) for novel trials. [N=9 control mice, 7 Cre mice. pairwise t-tests p>0.05 (Holm corrected)] * indicates p<0.05, ** indicates p<0.01

We stereotactically injected *Stx3^fl/fl^* mice with a virus encoding the Ca^2+^-indicator GCaMP (AAV-hSyn-jGCaMP7f). In addition, mice were injected with AAVs encoding either Cre-recombinase with mCherry (AAV-CaMKIIa-mCherry-IRES-Cre, “Cre mice”), or just mCherry (AAV-hSyn-mCherry, “control mice”). To allow *in vivo* imaging of these neurons, we placed an imaging window over the left dorsal CA1 region. As a conservative measure of Cre recombination, we only analyzed the Ca^2+^-activity of mCherry-positive cells (Fig S3C, S8A-B).

In order to obtain robust recordings of a large population of CA1-region place cells while engaging psychological processes analogous to the freely moving, novel-arm T-maze behavior above (Fig 2), we trained head-fixed mice to perform a virtual reality novel-arm Y-maze task (Figs 3B& S9A, <u>Movie S1</u>). In the novel-arm T-Maze described above (Fig 2E), Cre-injected and control mice differed significantly in their occupancy of the arms during test. This difference in novel arm preference, however, would complicate the comparison of neural activity across groups. Therefore, instead of allowing mice to choose which arm to explore, we controlled the occupancy of the different arms of the virtual Y-maze by forcing mice to take either a left or right “turn” on each trial as they ran forward on a fixed axis treadmill. During training, we forced exploration of 1 of the 2 virtual arms (“familiar”) for 5 blocks of trials. To assess the effects of novel experience on neural coding while ensuring sufficient exploration of both the familiar and novel arms during the test block, we randomly interleaved trials in which the mouse was forced to run down either the “familiar” or the “novel” virtual arm. We placed un-signaled reward zones in distinct locations on each arm (Fig 3B, shaded regions) to detect whether mice could distinguish the two arms of the maze. As mice learn the locations of the rewards, they should show distinct reward approach behaviors (e.g. slowing and licking at different locations) on the two arms. This was repeated for 5 days, and on day 6, mice ran interleaved trials on both arms for the whole session during both training and testing – essentially turning both arms into a “familiar” arm as a within-animal control.

Both Cre-injected and control mice rapidly learned the context-dependent reward locations and reliably distinguished between the two arms (Fig 3C & D, S9B-F). Despite comparable licking accuracy, control mice licked at a higher rate as they approached the reward zone for both familiar and novel arms. This difference suggests that abolishing LTP may weaken predictive reward coding while sparing the general ability to navigate towards rewards.

Strikingly, both groups had a large population of place cells that were stable in their spatial representation across days and displayed comparable remapping between familiar and novel arms (Fig 3E-J, S10). Additionally, there was no difference between groups in the fraction of cells that are place cells, the stability of spatial representations within a day, place cell remapping across arms, or the ability to decode position from neural activity (Fig S10 & S11). Place cells in the CA3 region and dentate gyrus of the hippocampus were also intact in the subset of Cre-injected mice for which we could image GCaMP activity in these regions (Fig S12). In addition, both groups showed indistinguishable remapping patterns on a separate context-discrimination task in which remapping patterns in ambiguous environments depend on previous experience (*50*) (Fig S13). These results show that, surprisingly, spatial and contextual encoding in CA1-region neurons is grossly unchanged after eliminating CA1 LTP, giving insight into why the freely moving spatial learning behaviors were largely normal in mice lacking CA1 LTP.

To understand why spatial and contextual encoding may appear grossly normal in CA1 place cells in the absence of LTP, we built a simple computational model for passively inheriting versus actively learning a place cell representation from presynaptic CA3 inputs (Fig S14). While previous work suggests that NMDA-receptors are necessary to form stable and accurate place codes (*3, 7*), this model predicts that stable place representations can emerge in a sparsely active CA1 neural cell population without LTP by simply reading out upstream spatially selective inputs. Furthermore, this model correctly predicts differences between control and Cre-injected mice in several subtle place field properties for which we would not otherwise have an interpretation. Place fields from Cre-injected mice were narrower than those from control mice, and place cells from Cre-injected mice had more place fields on average than place cells from control mice (Fig S14E-H). Our model-derived hypothesis that a CA1 place cell can inherit spatial selectivity from upstream inputs without LTP is consistent with previous findings: place cells receive more presynaptic inputs inside their naturally occurring place fields (*51*), and increasing the excitability of a cell can unveil “hidden” place fields (*52*). Although CA1 pyramidal cells can become a place cell at any location (*53*), CA1-region LTP may most often function to stabilize or exaggerate pre-existing biases in connectivity. Beyond the CA1-region, this model suggests that sparse representations can pass through downstream brain regions intact even when connectivity is random.

As an extension of this hypothesis, we propose that control and Cre-injected mice may differ in additional spatial and contextual coding properties that are also known to differ between the CA3 and CA1 regions, with CA1 neural activity from Cre-injected animals appearing more CA3-like. For example, a greater proportion of place cells are recruited to represent goal/reward locations in the CA1 region but not the CA3 region, leading to a population “overrepresentation” in the CA1 region (*54*). Correspondingly, we found that control animals allocate a greater proportion of place cells to rewarded locations than Cre-injected animals (Fig 4A-B, S15). A recent study showed that optogenetically stimulating these reward coding place cells drives increased licking in a similar VR task (*49*), thus the difference in reward representation we observe may provide a neural substrate for the matching difference in anticipatory licking in this task (Fig 3C & D). In addition, control animals had a larger proportion of “reward cells” (*55*) that represented the approach to reward locations on both arms (Fig S15C & D), suggesting LTP is important for forming functional ensembles of cells that encode salient aspects of the environment (*56*). To test that the LTP-dependent overrepresentation of rewards strengthens spatial reward associations we reversed the reward locations on the arms of the Y-Maze on days 7 and 8 in a subset of mice (Fig S16). Both groups of animals quickly learned to lick at the new reward locations, but control mice took more trials than Cre mice to extinguish licking at the original reward zone.

**Fig. 4.**
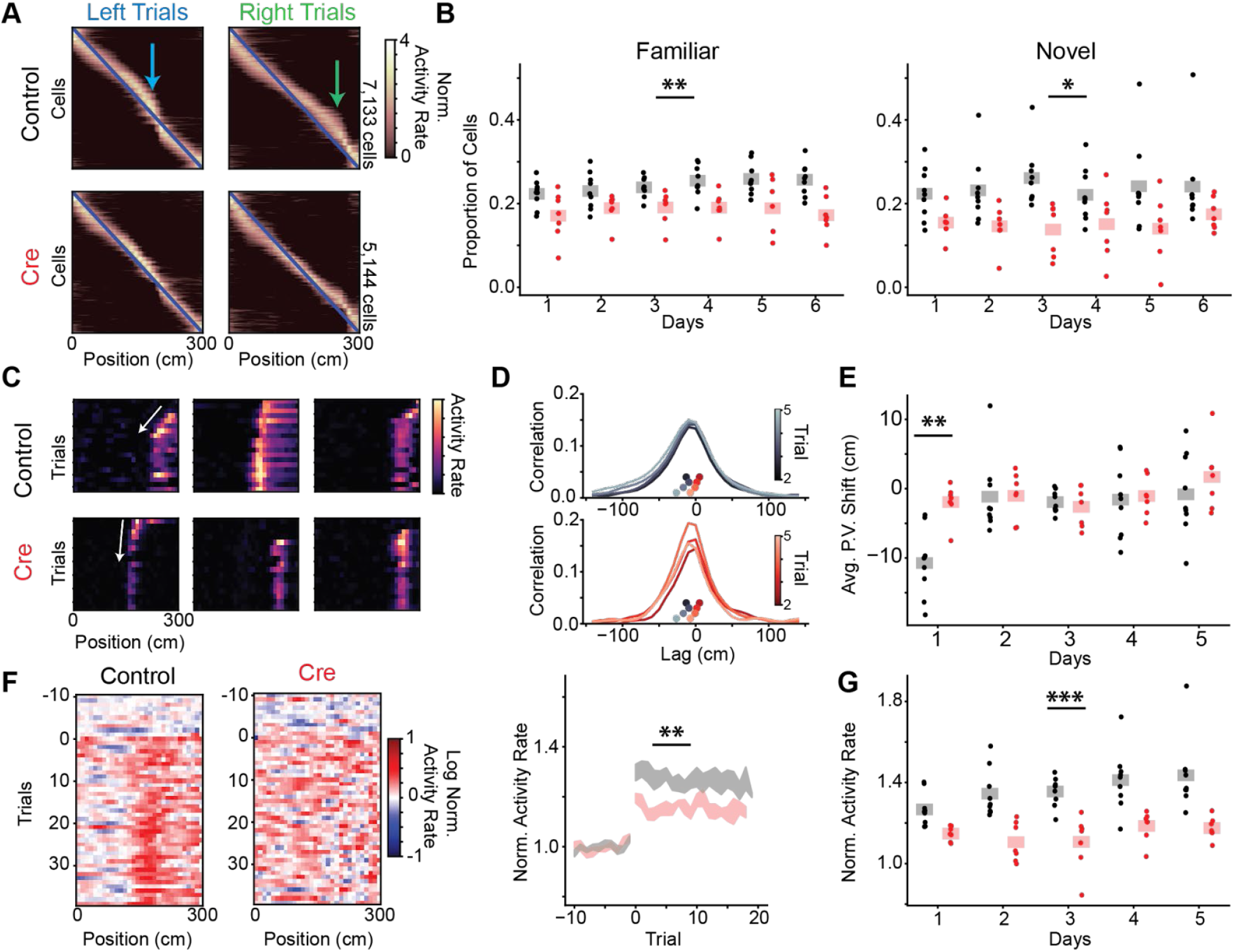
CA1-region LTP is required for both place cell overrepresentation of rewarded locations and for population novelty responses. **A-B**) LTP is necessary for recruitment of place cells to reward locations. **A**) Day 6 trial-averaged place cell activity (even trials only, odd trial sorted, z-scored) from all animals. Arrows indicate place cell overrepresentation of rewarded locations. Blue line shows a diagonal reference. **B**) Fraction of place cells with their peak near the reward zone for each animal on each day (left - familiar trials, right - novel trials) [N=9 control micel mice, 7 Cre mice, 6 days. Familiar trials mixed effects ANOVA: N=9 control mice mice, 7 Cre mice, 6 days, virus main effect p=0.003, day main effect p=0.068, interaction p=0.265. Novel trials mixed effects ANOVA: virus main effect p=0.017, day main effect p=0.653, interaction p=0.346] **C-E**) LTP is necessary for backward shifting of place cells in novel environments. **C**) *Top -* Example co-recorded place cells from a control mouse during novel arm trials on day 1. Cells display a backward shift of their place field during the initial trials (highlighted by white arrow). *Bottom -* Example co-recorded place cells from a Cre mouse during novel trials on day 1, demonstrating a less dramatic backward shift. **D**) Cross-correlation of population vector activity quantifies population backward shift. For illustration, we show across animal average population vector (PV) spatial cross-correlation between the first novel trial and the next 4 novel trials on day 1 (“initial novel trials”, top - control, bottom - Cre). Inset scatterplot shows the center of mass of the cross-correlation (“PV shift”) for each trial for both control (greyscale colormap) and Cre (red colormap). Control PV shifts are to the left of the Cre PV shifts indicating a larger magnitude backward shift for control trials. **E**) Average initial novel trial PV shift for each mouse on each day. Dots are the average PV shift for all pairs of initial novel trials on each day for each mouse (see Fig S17 for schematic). [N=9 control mice, 7 Cre mice, 5 days (day 6 excluded for different trial structure). Mixed effects ANOVA: virus main effect p=0.024, day main effect p=3.85×10^−4^, interaction p=0.028. Posthoc t-test: control vs Cre for day 1 p=0.002 (Holm corrected), all other days p>0.05] **F-G**) LTP is required for novelty-induced enhancement of the CA1 activity rate. **F**) *Left -* Population averaged log-transformed relative activity rate of trials in the final block on day 1 for an example control mouse. Trial 0 indicates the start of the final block. Each cell’s activity is normalized by it’s average activity in the 10 trials before the start of the final trial block. *Middle* - Cre mice display a smaller increase in activity for novel environments. Same as (left) for an example Cre mouse. *Right -* Averaged relative activity rate on each novel trial for day 1. Data are shown as across animal mean SEM. [N=9 control mice, 7 Cre mice. Posthoc t-test (see G) p=0.002 (Holm corrected)]. **G**) The vertical axis represents the mean relative activity rate across novel trials for each mouse on each day. [N=9 control mice, 7 Cre mice, 5 days (day 6 excluded for different trial structure). Mixed effects ANOVA: virus main effect p=4.59×10^−5^, day main effect p=0.005, interaction p=0.208] * indicates p<0.05, ** indicates p<0.01, *** indicates p<0.001

Beyond reward coding, neurons from the CA3 and CA1 regions differ in their responses to novel environments in two key respects. First, CA1-region place cells tend to shift their firing fields backwards (i.e. opposite the direction of travel) in novel linear environments (*57*). The magnitude of this shift is greater in CA1-region neurons than CA3-region neurons (*58*). This backward shift is thought to rely on LTP and be important for forming a predictive code for space in the CA1 region (*59, 60*). Indeed, we also found that LTP is necessary for the backward shift of spatial representations normally seen in a novel environment (day 1, Figs 4C-E, S17). We quantified this shift at the neural population level by calculating the center of mass of the spatial cross-correlation of population vectors on pairs of novel arm trials (Fig 4D) (*61*). Second, CA1-region neurons but not CA3-region neurons nonspecifically increase their firing rate in novel environments (*62, 63*). Confirming these previous reports, exposure to novel environments caused an increase in activity rate for control mice (Fig 4F, S18). The increase in activity for Cre mice was significantly smaller (Fig 4F &G), indicating that CA1 LTP is required for novelty modulation of activity rate. Unlike the backward shifting of place fields, we observed this phenomenon on each day of the experiment.

Collectively, the unimpaired contextual and spatial coding with the absence of typical novelty responses in Cre-injected mice provide a physiological basis for understanding the observed changes in the hippocampus-dependent behaviors of freely moving animals (Fig 2). Furthermore, the averaged population activity rate (Fig 4F) visually resembles the potentiation seen in LTP slice recordings. The *Stx3* deletion abrogates LTP (Fig 1C), diminishes the sustained increase of neural activity after novel context exposure (Fig 4F&G), and eliminates novel context-induced exploration/learning (Fig 2E). These data demonstrate that CA1-region LTP likely mediates a specific novelty computation and reinforces the utility of linking cellular processes to spatial computations in order to understand their roles in learning/behavior.

Here, we demonstrated a new approach for precisely probing the contribution of LTP to neural coding algorithms and behavior without interfering with basal synaptic transmission. We show that *Stx3* is essential for LTP induction in the hippocampus and striatum across a range of experimental variables, indicating a universal role for *Stx3* in LTP. Isolating LTP in this fashion revealed the surprising result that many behaviors known to require neuronal activity and NMDA-receptor function in the hippocampus or striatum do not require LTP within those regions. To understand the disconnect between the large physiological effects and the isolated behavioral changes induced by the *Stx3* deletion, we chose the CA1-region as an exemplar for investigating what LTP can contribute to neural coding within a single brain region. The results show that, rather than driving the formation of population codes necessary for expression of many forms of memory, LTP provides a mechanism to incorporate saliency signals into existing neural representations.

## Acknowledgements

We thank D. Luna, G. Wang, L. Ho, A. Diaz, and M. Sosa for help with animal care and training.

## Author Contributions

Conceptualization: KK, MHP, TCS

Methodology: KK, MHP, TCS

Investigation: KK, MHP, RY, RS

Visualization: KK, MHP

Funding acquisition: TCS, JBD, LMG

Project administration: KK, MHP, JBD, LMG, TCS

Supervision: JBD, LMG, TCS

Writing – original draft: KK, MHP, JBD, LMG, TCS

Writing – review & editing: KK, MHP, RY, RS, JBD, LMG, TCS

## Data and materials availability

All two-photon data will be made available on the DANDI repository upon acceptance. All two-photon analysis code will be made available upon acceptance.

## Supplementary Materials for

**This PDF file includes:**

Materials and Methods
Supplementary References
Supplementary Text
Figs. S1 to S18

**Other Supplementary Materials for this manuscript include the following:**

Table S1
Movie S1

### Materials and Methods

#### Subjects

All procedures were approved by the Institutional Animal Care and Use Committee at Stanford University School of Medicine. For all hippocampal experiments, we used male *Stx3^fl/fl^* mice (*65*) due to the known effects of estrous cycle on hippocampal LTP. For D1-Stx3 experiments, *Stx3^fl/fl^* mice were bred into a background carrying D1-tdTomato^+/-^ [JAX016204, (*66*)] and D1-Cre^+/-^ [STOCKTg(Drd1-cre)EY262Gsat/Mmucd, (*67*)]. Only mice heterozygous for D1-Cre+/- and D1-tdTomato^+/-^ were used. Male and female adult mice were used for D1-Stx3 experiments. All mice were housed in groups of between two and five same-sex littermates. After surgical implantation or virus injection, mice were housed in transparent cages and kept on a 12-hour light/dark schedule, and mice allocated to head-fixed behavior were housed with a running wheel from ∼1month of age. All experiments except DMP Barnes, Pavlovian conditioning, and self-administration were conducted during the light phase. Mice were between 5-16 weeks old for slice physiology experiments, 2-4 months at the time of virus injection for freely moving behavior experiments, 2-6 months at the time of D1-Stx3 behavior, and between 3 and 5 months at the time of surgery for two photon experiments.

#### Statistics

All numerical data in figures are means ± SEM. Whenever possible, the experiments were done in a blind manner. We followed guidelines for the proper use of parametric/linear models (*68*). A parametric model was considered proper if the residuals followed a normal distribution; if residuals did not lie on the identity line of the Normal QQ plot, and the data were lognormally distributed (according to the Shapiro-Wilk test) then the data were transformed by natural log and subsequent parametric analysis was run. If the data were neither normally nor lognormally distributed then a non-parametric test was used. Statistics on the electrophysiology and freely moving behavior were run using Prism 9 (GraphPad). When appropriate, repeated measures ANOVA was used with the Geisser-Greenhouse’s epsilon correction for sphericity and Holm-Šídák corrected multiple comparisons. ANOVAs were two-way unless stated otherwise. T-tests were always two-tailed. Analyses of virtual reality behavior and Ca^2+^ imaging data: Linear mixed effects models were run using the statsmodels package (https://www.statsmodels.org/stable/index.html). Mixed effects ANOVAs and subsequent posthoc t-tests were run using the pingouin package (https://pingouin-stats.org/) with a Holm step-down Bonferonni correction for multiple comparisons. For the two photon imaging data, distributions were assumed to be normal but this was not formally tested. No statistical methods were used to pre-determine the number of mice to include in this study, but our sample sizes are similar to those reported in previous publications.

#### Stereotaxic injection for electrophysiology or freely-moving behaviors

AAVs (serotype DJ, produced by HHMI Janelia Farms) expressing Synapsin-EGFP-CRE and Synapsin-EGFP-deltaCRE/ΔCre were injected into *Stx3^fl/fl^* mice. Mice were anesthetized with tribromoethanol (250 mg/kg, both viral conditions for open field, elevated plus maze, delay fear conditioning) or ketamine (100 mg/kg)/xylazine (10 mg/kg) (electrophysiology, T-Mazes, DMP Barnes, and trace fear conditioning) and head-fixed with a stereotaxic device. A glass micropipette with a long, narrow tip was pulled using a micropipette puller to deliver the virus. The glass pipette was slowly lowered to the target area and left for 2 min before virus injection. Virus solution was injected at an infusion rate of 100 nl/min and withdrawn 5 min after the end of injection. 300 nL of virus was infused into 4 sites – bilaterally in 2 different anterior-posterior positions (AP: -2 and -2.5, ML: 1.42, DV: -1.24). The mice were used for experiments at least 2 weeks after the virus infusion.

#### Brain slice preparation

Transverse hippocampal slices were prepared from *Stx3^fl/fl^* mice 2-3 weeks after virus injection using standard techniques (*69*). Mice were anesthetized with isoflurane, decapitated, and the brain was briefly exposed to chilled “slicing” solution containing 75 mM Sucrose, 87 mM NaCl, 2.5 mM KCl, 0.5 mM CaCl2, 1.25 mM NaH2PO4, 7 mM MgCl2, 25 mM NaHCO3, 1 mM Na-Ascorbate, and 10 mM D-glucose. The hippocampus was isolated and placed against an agar block. Transverse slices (300 μm) were cut with a tissue vibratome (Leica VT1200 S, Germany) in the “slicing” solution. Slices were left to recover for 30 min at 32°C in “slicing” solution and then at room temperature for 1 hr in an interface chamber with “storage” solution containing 125 mM NaCl, 2.5 mM KCl, 1.25 mM NaH2PO4, 1 mM MgCl2, 1 mM CaCl2, 25 mM NaHCO3, and 15 mM D-glucose.

Oblique horizontal slices were obtained from *Stx3^+/+^ or ^fl/fl^* mice on a D1-cre^+/-^;D1-tdT^+/-^ background using standard techniques (*70*). Briefly, animals were anesthetized with isoflurane and decapitated. The brain was exposed and chilled with ice-cold artificial CSF (ACSF) containing 125 mM NaCl, 2.5 mM KCl, 2 mM CaCl2, 1.25 mM NaH2PO4, 1 mM MgCl2, 25 mM NaHCO3, and 15 mM D-glucose. Slices were left to recover in ACSF for 30 min at 34°C and then at room temperature for an additional 30 min before recording.

#### Electrophysiology recording

All recordings were conducted in a blind manner. The slices were transferred to the recording chamber and were continuously perfused at 30-32°C with ACSF. All solutions were saturated with 95% O2 and 5% CO2. ACSF osmolarity was 300∼310 mOsm and pH was 7.4.

##### Hippocampal recordings

Whole-cell voltage clamp recordings were made with borosilicate glass pipettes (2.5∼4 MΩ) filled with a Cs^+^-based low Cl-internal solution containing 126 mM CsMeSO3, 10 mM HEPES, 1 mM EGTA, 2 mM QX-314 chloride, 0.1 mM CaCl2, 4 mM MgATP, 0.3 mM Na3GTP, 8 mM Na2-phosphocreatine (280∼290 mOsm, pH 7.3 with CsOH). 0.5% biocytin was included in the internal solution to later visual morphology. 50 μM picrotoxin was included in the bath solution, and the CA3 region was cut to prevent over-excitability in the presence of picrotoxin. CA1 pyramidal neurons were visually identified by conventional IR-DIC optics, and only EGFP^+^ neurons identified by epifluorescence were recorded. AMPAR-EPSCs were evoked at 0.1 Hz by electrical stimulation of schaffer-collaterals using tungsten electrodes (Matrix electrode, 2 × 1, FHC Cat# MX21AEW(RT2)) positioned in the S. radiatum. LTP baseline recordings were performed in voltage clamp holding the cell at -70mV, and baseline stimulation intensity was adjusted to obtain 70-100pA responses with no synaptic failures. LTP was induced by 2 trains of high frequency stimulation (100 Hz, 1 s) separated by 20 s, while cells were depolarized to 0 mV. This induction protocol was applied within 11 min of achieving whole-cell configuration, to avoid “wash-out” of LTP (*71*). The access resistance was < 27MΩ for LTP recordings and recordings were excluded if the access resistance changed >25%. The AMPAR/NMDAR ratio was calculated as the peak averaged AMPAR EPSC at -70 mV divided by the averaged NMDAR EPSC measured 50 ms after the onset of the dual component EPSC at +40 mV. Paired pulse ratio (PPR) was calculated as the ratio of the peaks of the 2nd EPSC/1st EPSC. Access resistance was < 20MΩ for basal transmission recordings and were discarded if resistance changed >20%.

##### Striatal recordings

Recordings were only performed from D1-tdTomato^+^ neurons. miniature Excitatory/Inhibitory PostSynaptic Currents (mEPSC/mIPSC) were recorded in the presence of TTX (0.5 µM) in the bath using whole-cell voltage-clamp with the Cs^+^-based low Cl^−^ internal solution described above. mEPSCs were recorded while holding the neuron at -70mV and mIPSCs at +8mV (liquid junction potential not corrected). The access resistance was < 20MΩ and recordings were excluded if resistance changed > 20%. The perforated patch and LTP induction protocol was adapted from a previous study (*23*). Electrical access was achieved through the perforated-patch method using Gramicidin A (Sigma). Perforated patch was performed with a borosilicate glass microelectrode (3-3.5 MΩ), front-filled with 1 µl K^+^-based internal solution (135 mM KMeSO3, 8.1 mM KCl, 10 mM HEPES, 8 mM Na2-Phosphocreatine, 0.3 mM GTP-Na, 4 mM ATP-Mg, 0.1 mM CaCl2, 1 mM EGTA; pH 7.2-7.3; osmolarity 285-290 mOsm), and back-filled with 10 µl Gramicidin A-containing internal solution. The Gramicidin A-containing internal solution was made fresh before use: a stock solution of Gramicidin A (20 mg/mL) in dimethyl sulfoxide (DMSO) was prepared and diluted in the K^+^-based internal solution yielding a final concentration of 200 µg/mL. The fluorescent dye Alexa488 (10 µM) was also added in the internal solution to visualize the integrity of the perforated patch configuration throughout the recording. After the microelectrode formed a gigaseal with the cell membrane, access resistance was continuously monitored during perforation by applying a -5 mV pulse from a holding potential of -70 mV, under the voltage-clamp mode. A stable perforated patch normally formed within 30-60 minutes, with a stable access resistance of ∼30-50 MΩ. Then the recording was switched to the current clamp mode, and serial resistance was compensated with the amplifier bridge balance. Presynaptic inputs, recorded as EPSPs, were evoked by focal extracellular stimulation with a small theta glass electrode positioned 50∼100 µm from the recorded cell body. Stimulation intensity (0.2 ms, 5-30 µA) was adjusted to evoke stable EPSPs with an amplitude of around 2∼5 mV. EPSPs were evoked every 20 s for 5 minutes as the baseline. Then LTP was induced using the spike-timing-dependent plasticity (STDP) protocol. The protocol consisted of 15 trains of five bursts repeated at 0.1 Hz. Each burst consisted of three postsynaptic action potentials preceded by three presynaptic inputs (EPSPs) at 50 Hz (+5 ms). The postsynaptic spikes were evoked by direct somatic current injections (5 ms, 1-1.5 nA). During the induction, the postsynaptic cell membrane potential was depolarized to -70 mV. After the STDP induction, EPSPs were recorded for another 30 minutes to monitor the change of amplitudes. 100 µM picrotoxin was applied throughout the recording.

Recordings were performed using a Multiclamp 700B (Molecular Devices) and monitored with WinWCP (Strathclyde Electrophysiology Software). Signals were filtered at 2 kHz and digitized at 10 kHz (NI PCIe-6259, National Instruments). Offline analysis used Clampfit 10.0 (Molecular Devices), MiniAnalysis (Synaptosoft), Easy Electrophysiology (RRID:SCR_021190), and custom MATLAB (Mathworks) software.

#### Behavior

All mice were handled for at least 3 days (>5min/day) before behavioral experiments. All video tracking experiments except fear conditioning were analyzed with the Viewer III tracking system (Biobserve) followed by custom MATLAB scripts.

##### Open field

Mice were individually placed into the center of the open field chambers (34cm × 34cm × 40 cm, white) and allowed to freely explore for 30 min. The total traveling distance and time spent in the center area were analyzed.

##### Elevated plus maze

The gray maze had four 30cm × 8 cm arms. Two “open” arms did not have walls, while 10 cm high walls enclosed the two “closed” arms. The maze was elevated 40 cm over the floor. At the beginning of the test, each mouse was individually placed at the junction of an open and a closed arm, facing the open arm, and was allowed to freely explore for 5 min. The time spent in the open versus the closed arms was analyzed.

##### Delay fear conditioning

The fear conditioning chambers (Coulbourn Instruments) were located in the center of a sound attenuating cubicle and were equipped with overhead infrared cameras to record behavior. A ventilation fan provided a background noise at ∼55 dB, and pans containing each respective odorant solution were placed below the floor. The conditioning chamber had stainless steel rods connected to a shock source, was cleaned with 10% ethanol, and had 2 metal and 2 transparent walls. After a 2 min baseline period, the mice received three tone (30s, 85 dB, 2 kHz)-footshock (0.75mA, 2s duration, delivered immediately after tone) pairings separated with a 1 min inter-trial-interval (ITI). The footshock co-terminated with the tone. The mice were returned to their home cages after 60s. In the context test 24hr later, mice were placed back into the original conditioning chamber for 5 min. 48hr after conditioning, mice were placed into an altered context: the floor was a plastic sheet with paisley design, all 4 walls were decorated with high contrast print, and the odor was 1% vanilla. After a 5 min baseline period to measure altered-context freezing, a tone (85 dB, 2 kHz, 1min) was delivered to measure cue-induced freezing. The context and altered context tests were repeated 7 and 8 days after conditioning, respectively. The mice were recorded with the FreezeFrame software and analyzed with the FreezeView software (Coulbourn Instruments). Motionless bouts lasting more than 1s were classified as freezing. During training, the freezing percentages in the 2 min exploration period, the 1 min period after each foot shock, and the tone were summarized as an indication of fear memory acquisition. The same mice were used in the following order: open field, elevated plus maze, and delay fear conditioning.

##### Trace fear conditioning

This procedure was adapted from (*25*). The apparatus and contextual cues were identical to above with the following procedural differences: mice received five tone (30s, 70dB, 2 kHz)-footshock (0.5mA, 2s) pairings separated by a 60s-120s ITI. Critically, a 15s “trace” period was inserted between the end of each tone and the start of the footshock. Trace conditioning using this interval is GluR1-and hippocampal-dependent (*25, 72*). 24hr after conditioning, the mice were placed in the altered context – after a 180s baseline period, the tone was presented for 180s. 10 days after conditioning, the mice were placed in the original conditioning chamber for 5 minutes. Hippocampal lesions disrupt expression of contextual fear memory at this time point (*27*).

##### Delayed Match to Place (DMP) Barnes Maze

DMP Barnes was performed during the mice’s dark phase – the only illumination before the session began was the experimenter’s red-light headlamp. An air filter was turned on for background noise (57dB). The maze consisted of a raised circular open platform (92cm diameter) with 20 equally spaced holes (hole diameter, 5cm) along the perimeter. An escape box (7cm deep, 7cm width, and 10cm length) was placed underneath the designated target hole, and false escape boxes were placed under the remaining holes, made of the same color/texture material as the escape box. Extra-maze cues were placed on the surrounding walls and remained stationary throughout the procedure. The mice were habituated to the goal box and scent of 1% Virkon-S cleaning solution for 1min ∼24hrs before beginning training. To begin the session, a mouse was placed in the center of the maze in a holding chamber (15cm × 15cm) for 10s. Then a bright light was turned on as the chamber was lifted – beginning the session. The light remained illuminated until the mouse entered the correct escape port – concluding the session. If the mouse did not enter the target escape hole within 90s, the mouse was gently guided to the correct hole and the light turned off. The mouse was allowed 10s in the goal box before being returned to its home cage. After each trial, the maze was cleaned with Virkon disinfectant to scramble olfactory cues. Each day consisted of 4 trials with a 20min ITI, an ITI that exposes a deficit in the DMP Morris water maze with hippocampal lesions or NMDAR antagonist (*28*). The goal box was moved every day to a cardinal point in the same order for all mice (N, W, E, S). X,Y coordinates of the center of the mouse were recorded over time. The escape zone was defined as the area including the target hole and its two neighbors, with 2.5cm dilation.

##### Food T-Maze

The dimensions of each transparent arm were 25cm×8cm×20cm (L×W×H) with horizontal or vertical gratings differentiating the left vs right arm walls. Mice were single housed and maintained at 85-90% body weight with *ad libitum* access to water. Mice were given 10 reward pellets (20mg palatable food pellet: 27% Fat, Bioserve catalog # F06649) in their homecage 24hrs before conditioning to avoid neophobia. To begin, a mouse was placed in the start arm and was allowed access to all arms of the maze during training and test. Training consisted of five, 2min sessions separated by a 1min ITI. 1 reward pellet was placed on a metal mesh receptacle at the end of the rewarded arm (within 6cm of the end of the arm, “rewarded” side counterbalanced across groups). The “scent control” arm had the same metal mesh receptacle, except the food pellet was inside the mesh and inaccessible to the mouse – avoiding the confound of scent and sight of the pellet/receptacle. The test session was the empty T-Maze (all receptacles were removed - pilot experiments demonstrated mice chew the receptacles when empty). There was a 1min ITI between training and test, and the test session began when all 4 paws left the start arm. The following day, the mice were allowed access to the same maze to test a 24hr memory recall for 2 trials. Test performance was measured by the difference score: (time spent in the reward arm) – (time spent in the scent control arm).

##### Novelty T-Maze

Adapted from (*31*). The dimensions of each transparent arm were 30cm×10cm×20cm (L×W×H). This is a slightly larger T-Maze than the one used for the food T-Maze and was constructed of different plastic. The two mazes also differed by the time of day, extra-maze visual/auditory cues, and method of transport. During training, the entrance to one arm (“Novel arm”, assignment counterbalanced across groups) was blocked with black, matte plastic. Training consisted of five, 2min sessions separated by an ITI in which each subject was allowed to explore the start and “familiar” arms. Following an ITI, each mouse was allowed to explore all arms of the maze – the novelty preference test. This session was 2min and began when all 4 paws left the start arm. The novelty T-Maze had 2 different ITI conditions – 1min and 24hr. This ITI was consistent for both training and test (i.e. 1min between each training session and 1min between the last training session and the test). All mice experienced both ITI conditions in contexts differentiated by background noise, extra-maze visual cues, time of day, method of transport. The test order was counterbalanced across groups. Test performance was measured by the difference score: (time spent in the novel arm) – (time spent in the familiar arm). A subset of mice went through the novelty T-maze, DMP Barnes, and Trace Fear Conditioning (in that order). Another subset only went through the novelty T-maze and reward T-maze.

##### Cocaine locomotor sensitization

Open field chambers were 25.4cmx25.4cmx25.4cm. Mice were habituated to intraperitoneal injections for 2 days in their homecages before testing. Mice received a (10µL solution)/(g body weight) intraperitoneal injection immediately before each of the four 1hr sessions. On days “-2” and “-1,” mice received saline. On day “0” mice received 20mg/kg cocaine (Sigma, CAS 53-21-4), and 1 week later received the same concentration.

##### Rotarod

An accelerating rotarod for mice (IITC Life Science) was used. Mice received 3 trials/day over 4 days. The rotarod was activated after placing mice on the motionless rod. The rod accelerated from 4 to 40 revolutions per minute (days1-3) or 8 to 80 rpm (day4) - both over the course of 5 minutes. Each trial ended when a mouse fell off, made 1 complete revolution while hanging on, or reached 300 seconds.

Pavlovian conditioning and operant self-administration occurred during the dark cycle (mice were in a reverse light-cycle room). Mice were single-housed and maintained at 85-90% starting 3 days before the first day of Pavlovian conditioning and were maintained on food restriction until the first habituation day for self-administration, after which they were fed *ad libitum*. Mice received an additional 5 food pellets in their homecage (20mg palatable food pellets, 27% Fat, Bioserve catalog # F06649) to prevent neophobia during training. We used standard mouse operant chambers (Med Associates) individually enclosed in ventilated sound attenuating chambers and initiation of the session illuminated a red house light.

##### Pavlovian appetitive conditioning

Mice were given two sessions in which they were habituated to the operant chambers. During these sessions, the house light was illuminated and one 20mg palatable food pellet was delivered once a minute into the food receptacle. Subsequently, pavlovian conditioning was initiated – the mice received a 10 second compound conditioned stimulus (CS, cue light and tone) on a variable interval 60s (VI-60) schedule. The food pellet was delivered halfway into the CS (after 5 seconds) and beam breaks into the food port were recorded. 14 CS presentations occurred per session over 9 days. Learning was assessed by a conditioned approach (CA) score: the head entry rate during the CS was subtracted from the head entry rate during the inter-trial interval (*42*).

##### Operant self-administration

After the final pavlovian conditioning session, the mice underwent a 1 hr conditioned reinforcement test (*73*). The nose poke ports were uncovered in the same chambers, and the mice were allowed to nose poke in the illuminated, active port for a 2 s presentation of the same compound cue used in pavlovian conditioning, but no food. To prime the mice to respond, 1 food pellet was placed at the entrance of the active, illuminated port, and nose pokes in the inactive port had no programmed consequences for all testing described here. The day after the conditioned reinforcement test, we began 2 days of habituation to the food self-administration procedure. Initiation of the session illuminated the active nose poke port, and we allowed food restricted mice to self-administer the 2s CS and 1 food pellet on a fixed-ratio-1 (FR-1), 5-second timeout schedule i.e. one nose poke in the active port delivered 1 food pellet. Then food restriction was removed for the remainder of testing. In order to study food self-administration without confounds of the pavlovian CS, we removed the CS for FR-1 and progressive ratio testing. After 3 days at FR-1, the mice underwent 4 days of progressive ratio (PR-3) testing. Here, the response requirement increased by 3 for each successive reinforcer (1+4+7 nose-pokes required to receive 3 food pellets, etc). The final breakpoint is defined as the last completed response requirement within the 1 hr session.

#### Histology/Imaging

Mice were deeply anesthetized with tribromoethanol and transcardially perfused with PBS followed by 4% PFA. Brains were removed and post-fixed in 4% PFA for 24hrs (1hr if proceeding with X-gal staining) and then transferred to 30% sucrose in PBS for 48hrs at 4°C. 100μm sections were taken on a cryostat (CM3050 S; Leica Biosystems) and mounted on glass slides with DAPI Fluoromount-G (Southern Biotech). Images were collected using a 10x objective on a VS120 or VS200 slide-scanning microscope (Olympus). We annotated standard Paxinos plates (*74*) with viral spread from each mouse we used in behavior.

##### X-gal staining

Slides with sections were rinsed in PBS 3 times, each for 5 min. Sections were then developed in a solution containing 5 mm K-ferricyanide, 5 mm K-ferrocyanide, 2mM MgCl_2_ and 1 mg/ml X-gal) at 37°C for 24 h in the dark. Finally, samples were rinsed with distilled water twice, slightly air-dried, and mounted with coverslips. Transmitted light imaging was then performed.

##### Dendritic morphology

slices used in electrophysiology were stained with streptavidin 555 to detect biocytin-filled cells for morphological analyses, and images were acquired using a Nikon A1 Eclipse Ti confocal microscope at 20× (branch order analysis) and 60× (dendritic spine density) objectives, operated by NIS-Elements AR acquisition software. The image stacks then went through a 3D deconvolution (NIS-Elements) using the known optical parameters. Neural morphology at 20x for branch order analysis was first analyzed using the TREES toolbox (*75*) and then fine-tuned with NeuronStudio (*76*). Dendritic spine density was quantified from 60x images in NeuronStudio. In each CA1 sub-region, we analyzed 10 dendritic segments of at least 10µm length each, all at least 100µm from soma (avoiding primary branches).

##### RNAscope

RNAscope *in situ* hybridization was performed using the manufacturer’s guidelines for fixed tissue (12µm tick sections). We used the Multiplex Kit v2 and Drd1, Drd2, and Custom Stx3 probes. Slides were mounted with gold antifade media (ThermoFisher). Multi-channel tiled images were obtained using a laser scanning confocal microscope (20x objective, LSM900, Zeiss). Linear unmixing was performed. Data were then analyzed with CellProfiler (*77*).

#### *In vivo* imaging virus injections and window implants

*Stx3^fl/fl^* mice were first anaesthetized by an intra-peritoneal injection of a ketamine/xylazine mixture (∼85 mg/kg). Animals were also subcutaneously administered 0.08 mg Dexamethasone and 0.2 mg Carprofen. After the first hour of the procedure, anesthesia was maintained with 0.5-2% isofluorane. For control mice (n=9 Ctrl1-9), adeno-associated virus containing GCaMP7f (AAV1-hSyn-jGCaMP7f, AddGene viral prep #104488-AAV1) and a second virus encoding the static red indicator mCherry (AAVDJ-CaMKII-mCherry, Stanford Gene Vector and Virus Core) were mixed and drawn into the same 36 gauge Hamilton syringe (World Precision Instruments; GCaMP titer - 1 x 10^12^, mCherry titer - 5 x 10^12^). For cre-injected mice (n=7, Cre1-7), the same GCaMP virus and a virus encoding Cre recombinase and mCherry (AAVDJ-CaMKII-mCherry-IRES-cre, Stanford Gene Vector and Virus Core) were mixed and drawn into the syringe (cre titer – 5-10 x 10^12^). For both groups of mice, virus was injected in two locations in the left dorsal CA1-region (injection 1: -2 AP, -1.4 ML, 1.35 DV; injection 2: -2.5 AP, -1.4 ML, 1.35 DV) and two locations in the right dorsal CA1-region (injection 1: -2 AP, -1.4 ML, 1.35 DV; injection 2: -2.5 AP, -1.4 ML, 1.35 DV). Each injection was 300 *η*L (50 *η*L/min) and the needle was left in place for 5 minutes following each injection to allow for virus dispersal. In an additional mouse (Cre-8), we performed identical dorsal CA1 injections of the cre virus without GCaMP and then injected the same GCaMP virus into the left dorsal CA3 (-1.5 AP, -1.6 ML, 1.98 DV, 500 *η*L at 50 *η*L/min).

To provide optical access to the CA1 pyramidal cell layer, after virus injections were complete, an imaging cannula was implanted over the left dorsal hippocampus as previously described (*50, 78*). Imaging cannulas consisted of a 1.3 mm length stainless steel cannula (3 mm outer diameter, McMaster) glued to a circular cover glass (Warner Instruments, #0 cover glass 3mm diameter; Norland Optics #81 adhesive). Excess glass overhanging the edge of the cannula was shaved off using a diamond tip file. A 3 mm diameter craniotomy was performed over the left posterior cortex (Cre1-7 and Ctrl1-9 centered at -2 mm AP, -1.8 mm ML, Cre8 centered at -1.5 mm AP, -2 ML). The dura was then gently removed and the overlying cortex was aspirated using a blunt aspiration needle under constant irrigation with ice cold sterile artificial cerebrospinal fluid (ACSF). Excessive bleeding was controlled using gel foam that had been torn into small pieces and soaked in sterile ACSF. Aspiration ceased when the fibers of the external capsule were clearly visible. Once bleeding had stopped, the imaging cannula was lowered into the craniotomy until the coverglass made light contact with the fibers of the external capsule. In order to make maximal contact with the hippocampus while minimizing distortion of the structure, the cannula was placed at approximately a 10 degree roll angle relative to the animal’s skull. The cannula was then held in place with cyanoacrylate adhesive. A thin layer of adhesive was also applied to the exposed skull. A number 11 scalpel was used to score the surface of the skull prior to the craniotomy so that the adhesive had a rougher surface on which to bind. A stainless steel headplate with a left offset 7 mm diameter beveled window was placed over the secured imaging cannula at a matching 10 degree angle, and cemented in place with Met-a-bond dental acrylic that had been dyed black using India ink.

At the end of the procedure, animals were administered 1 mL of saline and .2 mg of Baytril and placed on a warming blanket to recover. Animals were typically active within 20 min and were allowed to recover for several hours before being placed back in their home cage. Mice were monitored for the next several days and given additional Carprofen and Baytril if they showed signs of discomfort or infection. Mice were allowed to recover for 14 days before beginning water restriction and VR training.

#### Two photon (2P) imaging

To image the calcium activity of neurons, we used a resonant-galvo scanning two photon microscope (Neurolabware). 980 nm (Ctrl1-5, Cre1-7) or 920 nm (Ctrl6-9, Cre8) light (Coherent Discovery laser) was used for simultaneous stimulation of GCaMP and mCherry in CA1. 920 nm light was used in cases where mCherry fluorescence bleedthrough into the green photomultiplier tubes exceeded usable levels when stimulating at 980 nm. 920 nm light was also used to image CA2/3 and dentate gyrus (DG). Laser power was controlled using a pockels cell (Conoptics). Average power for excitation ranged from ∼35 mW to ∼80 mW measured at the front face of the objective (CA1 imaging: Nikon 16x, 3 mm WD, 0.8 NA; DG & CA3 imaging: Leica 25x, 3 mm WD, 1.0 NA). A 512 x 796 pixel field of view (FOV) was collected using unidirectional scanning at 15.46 Hz. Cells were imaged continuously under constant laser power during each block of trials (described below). To minimize photodamage and photobleaching, the pockels cell was used to reduce laser power to minimal levels between trial blocks.

Putative pyramidal cells were independently identified using the Suite2P software package (https://github.com/MouseLand/suite2p) on each session. Segmentations were curated by hand to remove ROIs that contained multiple somas, dendrites, or contained cells that did not display a visually obvious transient. Cells that did not contain a clear mCherry signal were also removed. The same FOV was imaged during each session. Custom code was used to find cells that were successfully tracked across imaging sessions. Baseline fluorescence was calculated for each cell within each block of trials independently using a sliding window maximin procedure (modification of default suite2p procedure). *Δ*F/F was then calculated for each cell as the change in fluorescence from baseline divided by baseline. These traces were median filtered to suppress optical shot noise and then deconvolved with a canonical calcium kernel to obtain “activity rates”. We do not interpret this result as a spike rate. Rather, we view it as a method to suppress the asymmetric smoothing on the calcium signal induced by the indicator kinetics.

#### Virtual reality (VR) design

All virtual reality environments were designed and implemented using the Unity game engine (https://unity.com/). Virtual environments were displayed on three 24 inch LCD monitors that surrounded the mouse and were placed at 90 degree angles relative to each other. A dedicated PC was used to control the virtual environments and behavioral data. This PC was synchronized with calcium imaging acquisition using TTL pulses sent to the scanning computer on every VR frame. Mice ran on a fixed axis foam cylinder and running activity was monitored using a high precision rotary encoder (Yumo). Separate Arduino Unos were used to monitor the rotary encoder and control the reward delivery system.

#### Water restriction and VR training

In order to incentivize mice to run, animals water intake was restricted. Water restriction was implemented 10-14 days after the imaging cannula implant procedure. Animals were given 0.8 - 1 mL of 5% sugar water each day until they reached ∼85% of their baseline weight and given enough water to maintain this weight.

Mice were handled for 3 days during initial water restriction and watered through a syringe by hand to acclimate them to the experimenter. On the fourth day, we began acclimating animals to head fixation (day 4: ∼30 minutes, day 5: ∼1 hour). After mice showed signs of being comfortable on the treadmill (walking forward and pausing to groom), we began to teach them to receive water from a “lickport”. The lickport consisted of a feeding tube (Kent Scientific) connected to a gravity fed water line with an in-line solenoid valve (Cole Palmer). The solenoid valve was controlled using a transistor circuit and an Arduino Uno. A wire was soldered to the feeding tube and capacitance of the feeding tube was sensed using an RC circuit and the Arduino capacitive sensing library. The metal headplate holder was grounded to the same capacitive-sensing circuit to improve signal to noise, and the capacitive sensor was calibrated to detect single licks. The water delivery system was calibrated to deliver ∼4 *μ*L of liquid per drop.

After mice were comfortable on the ball, we trained them to progressively run further distances on a VR training track in order to receive sugar water rewards. The training track was 450 cm long with black and white checkered walls. A pair of movable towers indicated the next reward location. At the beginning of training, this set of towers was placed 30 cm from the start of the track. If the mouse licked within 25 cm of the towers, it would receive a liquid reward. If the animal passed by the towers without licking, it would receive an automatic reward. After the reward was dispensed, the towers would move forward. If the mouse covered the distance from the start of the track (or the previous reward) to the current reward in under 20 seconds, the inter-reward distance would increase by 10 cm. If it took the animal longer than 30 seconds to cover the distance from the previous reward, the inter-reward distance would decrease by 10 cm. The minimum reward distance was set to 30 cm and the maximal reward distance was 450 cm. Once animals consistently ran 450 cm to get a reward within 20 seconds, the automatic reward was removed and mice had to lick within 25 cm of the reward towers in order to receive the reward. After the animals consistently requested rewards with licking, we began novel arm Y-maze training described below. Training took 2 - 4 weeks.

#### VR novel arm Y-maze

We sought to create a VR task that could probe the same phenomena as the freely moving T-maze behaviors in the other portions of the manuscript. However, by design, these tasks result in dramatic occupancy differences between the arms of the maze. This complicates statistical analyses of spatial and contextual coding. Furthermore, it is unknown if wildtype mice prefer to explore novel VR environments. To deal with this tradeoff we created a virtual Y-maze in which we forced mice to turn down different arms of the maze (Fig 3B, Fig S9A, <u>Movie S1</u>). Briefly, on each trial, the mouse began in a dark “timeout box” at the beginning of the track. The mouse self-initiated the trial by running forward on the ball. The animal then ran down the track to find a hidden reward and ran to the end of the track to end the trial. Whether the trial was a left or right turn was predetermined. The mouse was then teleported back to the “timeout box”, where it was forced to wait for a brief random duration (1-5 seconds, uniformly distributed) before beginning the next trial. This design did not allow us to assess the animal’s preference for each arm; however, it allowed us to test for the animal’s ability to distinguish the arms and examine novel context encoding while controlling for the occupancy of the different arms. The task essentially reduces to previously published VR novel context tasks (*79, 80*) while avoiding the sudden change in stimuli used in these tasks.

The animal needed to run 300 cm to complete each trial, but apparent motion along the track was a nonlinear function of movement on the ball. Smooth trajectories for left and right turns were created by fitting a cubic spline to a set of control points along the track. The left and right turn control points were symmetric, so that left and right turns were mirrored. The stem and arms of the maze were of equal length, but to create a smooth trajectory, the trajectories noticeably diverge approximately two thirds of the way down the stem of the Y-maze. Due to the uneven spacing of control points to create convincing trajectories, the visual speed as a function of movement on the ball slows near the turn and increases at the ends of the maze. As the spline trajectory parameter is linearly related to the animal’s motor action, we perform all spatially binned analyses as binned values of the spline parameter. These bins correspond to 10 cm of movement along the ball (30 bins to cover the track).

After initial running training, animals were trained to run in a “training Y-maze” to acclimate them to the spline trajectories. All walls of the training Y-maze were black and white checkerboards, with a black floor and a grey ceiling. The mice were required to lick to receive a hidden reward (i.e. no explicit visual cue marked the location of the reward) at the midpoint of either arm of the maze. Left and right turns were randomly interleaved on the training Y-maze. Once the animals ran smoothly and licked consistently to get the hidden rewards (1-3 days), we began the novel arm Y-maze task.

For the novel arm Y-maze, each section of the track had a dramatically different pattern of visual stimuli to aid the mice in distinguishing the arms (Fig S9A, <u>Movie S1</u>). The reward was omitted on 10% of trials. On each of the first 5 days of the experiment, the animals performed 6 blocks of trials. For the first 5 blocks of trials, animals were forced to go down only one arm (“familiar” arm) and the view of the other arm was blocked with a translucent virtual curtain (Fig S9A, <u>Movie S1</u>). In the sixth block, the virtual curtain was removed and familiar and novel arm trials were randomly interleaved. On the sixth day, the block structure was retained but familiar and novel trials were randomly interleaved from the first trial. Mice were randomly assigned to having the left or right arm as the familiar arm. In order to determine if the mice could distinguish the arms of the maze, they were required to lick at a hidden zone (25 cm wide) at the beginning of the left arm and the end of the right arm in order to receive a liquid reward. On the first block of trials on day 1, an automatic reward was dispensed at the end of the reward zone if the mouse did not lick within the reward zone. To mimic the freely moving task while getting sufficient trials to perform statistical comparisons, each of the first 5 blocks was a maximum of 5 minutes or 20 trials and the last block was 10 minutes or 40 trials. The inter-block interval was 1 minute.

On day 7, the procedure from day 6 was repeated for the first 2 blocks of trials. During the inter-block interval, we switched the reward locations on the left and right arms without changing any of the visual stimuli. An automatic reward was available on block 3 but removed for the remaining blocks. On day 8, the rewards remained in the reversed locations and left and right trials were randomly interleaved for all blocks.

#### Place cell identification and analyses

Place cells were identified independently on familiar and novel trials on each day using a previously published spatial information (SI) (*81*) metric 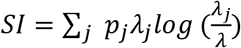. Where *λ_j_* is the average activity rate of a cell in position bin *j*, *λ* is the position-averaged activity rate of the cell and *p_j_* is the fractional occupancy of bin *j*.

To determine the significance of the SI value for a given cell, we created a null distribution for each cell independently using a shuffling procedure. On each shuffling iteration, we circularly permuted the time series of the cell relative to the position trace within each trial and recalculated the SI for the shuffled data. Shuffling was performed 1,000 times for each cell, and only cells that exceeded 95% of permutations were determined to be significant place cells.

Place cell sequences (Fig 3F&H, 4A) were calculated using a split halves procedure. The average firing rate maps from odd-numbered trials were used to identify the locations of peak activity and the activity on even-numbered trials is shown (z-scored across positions). When comparing place cell sequences for different trial types, we considered the union of place cells in both conditions.

#### Naïve Bayes decoding

To decode position from spatially binned population neural activity, we used the following decoder: 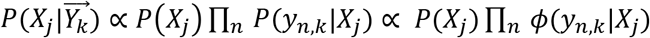, where 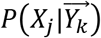 is the posterior probability that the mouse is in position *X_j_* given the position binned vector of neural activity rates 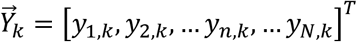 where *n* indexes the neuron identity and *k* indexes the unknown position. *P*(*X_j_*) is the prior probability the mouse is in position bin 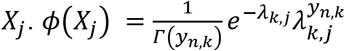, is a continuous quasi-Poisson potential function, where *λ_k,j_* is the trial-averaged activity rate of neuron *k* at position *j*. We assume that *P*(*X*) is uniform, reducing the model to a maximum likelihood decoder.

We used leave-one-trial out cross validation to assess the accuracy of the decoder. Error was calculated as the mean absolute deviation of the most likely position, 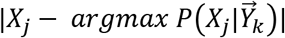. To compare errors across animals, we repeatedly randomly sampled an equal number of cells from each mouse with replacement to perform cross-validation (50 iterations per model size).

#### K-Winners-Take-All (KWTA) model for place cell inheritance

Our goal for this model was to write down the simplest network with plausible components that could simulate whether a CA1 population could inherit a place cell representation even without synaptic plasticity and, if so, how plasticity could change this representation. This model has only three main components: 1) spatially selective input neurons, 2) Hebbian plasticity on input weights by output neurons and, 3) recurrent inhibition between output neurons.

We considered a two layer neural network with a set of spatially selective input neurons 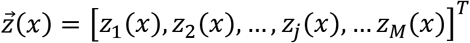, a set of output neurons, 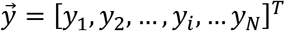, and a connectivity matrix, *W*, where *W_ij_* is the weight from input neuron *j* to output neuron *i*. Input neurons have radial basis function tuning for position, 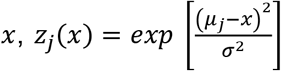, where *μ_j_* is the center of the radial basis function for neuron *j*. Basis functions were chosen to tile positions across the population, 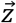. *σ*^2^ is the width of the radial basis function and is fixed for all input neurons. For Fig. S13, M=1000, N=1000, *σ*^2^ = 0.5, *x* ∈ [0,10].

We accomplished recurrent inhibition between output neurons using a K-Winners-Take-All approach. At every position along the track 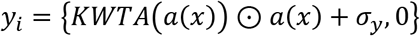, where 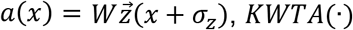 is a vector-valued function that applies the KWTA threshold and outputs a binary vector choosing the K winners, ⊙ denotes the elementwise product and *σ_y_* is additive noise to the output. *σ_z_* is a stimulus noise term. K=50, *σ_y_* ∼0.5*N*(0, 1), *σ_z_*∼0.05*N*(0, 1).

Weights were updated according a basic Hebbian learning rule at the end of each track traversal, *ΔW_ij_* = *ηz_j_y_i_*, where *η* is a constant. We required that weights had to be nonnegative. Weights also decayed to prevent explosive growth of synapse weight known to occur in Hebbian models, and additive Gaussian noise was included in the weight update. This gave the following weight update equation: 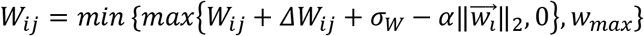. For “no LTP” model *η* = 0, *α* = 0 and for the “LTP” model *η* = 1 *x* 10^−4^, *α* = 9*x*10^−4^. For all simulations *σ_w_* = 0.001*N*(0, 1) and *w_max_* = 10.

We initialized weights using a lognormal distribution (mean=0, standard deviation = 0.5) (*82, 83*) though uniform weights also reproduced our results under different noise regimes (data not shown).

#### Novel arm activity rate increase

To assess whether overall neural activity rate increased in a block of trials while controlling for bleaching or within session FOV drift, each cell’s position-binned activity rate was divided by the mean activity rate of that cell in the 10 trials preceding the block of trials to be analyzed. We took the mean of this normalized activity rate across all cells to get a population normalized activity rate. This normalization gives a nonnegative value, where values less than one indicate a decrease in activity, and values greater than one indicate an increase in activity. For Fig. 4F & S18G, we took the log of this value for ease of visualization.

#### Place field statistics

##### Place field width

For cells with significant spatial information, field width was calculated as the spatial full width half maximum of the peak of the cell’s trial averaged activity rate map.

##### Number of fields per cell

Significant place fields were identified by a separate method independent of place cell identification. To identify significant place fields, we first calculate the trial averaged activity rate map for each cell. We then created a null distribution for these rate maps by randomly circularly permuting each individual trial’s rate map 1,000 times. Contiguous spatial bins with an activity rate that exceeded 99% of this null distribution were considered significant fields.

#### Population vector cross-correlation

Using a method previously described (*61*), we estimated trial x trial spatial shifts in population activity. For each pair of trials, we shifted one trial relative to the other (±300 cm, 10 cm spatial bins) and computed the population vector correlation at each lag (i.e. a spatial cross-correlation). We calculated the shift in population activity as the center of mass (COM) of this cross-correlation (Fig 4D-E & Fig S17). This gave an anti-symmetric matrix of COM values. To calculate the average initial shift of the population we averaged the upper triangular entries in the trial x trial matrix containing the first five novel arm trials.

#### Reward overrepresentation analysis

Place cells were considered to be reward zone cells if their location of peak activity was within the 50 cm preceding the front of the reward zone. Normalized population place cell activity was calculated by z-scoring each place cell’s trial averaged activity rate map and then averaging across all cells. To calculate the peri-reward zone population activity, we averaged this population metric within the same 50 cm preceding the reward zone.

#### Reward reversal analysis

Normalized lick rate in the previous reward zone was calculated as the average lick rate within a trial in 50 cm preceding the previous reward zone divided by the across trial averaged lick rate in the same spatial bins for day 7 blocks 1 and 2.

### Supplementary Text

#### Experience dependent hippocampal remapping

After completion of the novel arm Y-maze protocol, a subset of mice (Ctrl-1,3 & 5, Cre-1,2,4 & 5) were trained in the Rare Morph condition from (*50*). In this protocol, animals were trained to run in a set of virtual linear tracks with randomly placed cued rewards. The environments are distinct from the Y-Maze used in the rest of the two photon experiments. Further, these environments were designed to parametrically morph from one extreme to the other (Fig. S13A). A track was identified by its “morph value”, *S*, along this spectrum. The morph value can be translated into the horizontal frequency of sine waves on the wall, *f_h_*. On days 1-2 and 4-7 the mice only received randomly interleaved trials of the two extremes of the morph axis. On days 3 and 8-N, the animals received several “warmup” trials of randomly interleaved extreme morph values. During imaging, the animals saw randomly interleaved morph values from the full range of stimuli. After imaging was complete, animals continued trials in the extreme morph values to get the rest of their water for the day (Fig S13B). This Rare Morph condition was designed to give the animals a bimodal prior expectation over the frequency with which they should see these morphed environments (Fig. S13C).

In the original publication, we showed that remapping patterns in the intermediate environments are well explained by a model in which the hippocampus represents the posterior distribution over environments (Fig. S13D). This posterior estimate takes the following form: 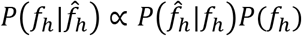, where *P*(*f_h_*) is the prior probability (Fig S13C), 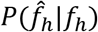 is the likelihood distribution which quantifies the noise in the sensory estimate of the stimulus (a Gaussian), and 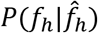 is the posterior distribution which is used for inference. As the animal had access to *f_h_* but not *S*, probabilistic inference was performed with *f_h_*. However, trials were evenly sampled on the *S* reference frame, so it was used for some analyses.

We can simulate a population of neurons that encodes this distribution exactly by positing that each neuron has some value of the inferred value of the stimulus, 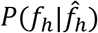, for which it prefers to fire, and that each neuron’s firing rate is proportional to the probability of the stimulus. This is equivalent to saying that each row in Fig. S13D is the activity rate of a neuron as a function of *S*. Calculating a trial x trial population cosine similarity matrix for this simulated population (Fig S13E) gives us a prediction of what we should see in the real data; trials should be essentially clustered into two groups (block diagonal structure of the similarity matrix). In both control and Cre animals, we found that this prediction holds, replicating the previous finding. Furthermore, as in our previous manuscript, the ideal posterior distribution can be reconstructed with high accuracy from the neural data alone for both control and Cre animals (Fig S13H). Thus, experience dependent remapping in the CA1-region does not require LTP. An experience dependent representation likely already exists in the CA3-region. A computational model from the original publication (21) suggests that such remapping does not require CA1 specific circuitry. This upstream representation can likely be inherited even without plasticity (Fig S14).

**Fig. S1.**
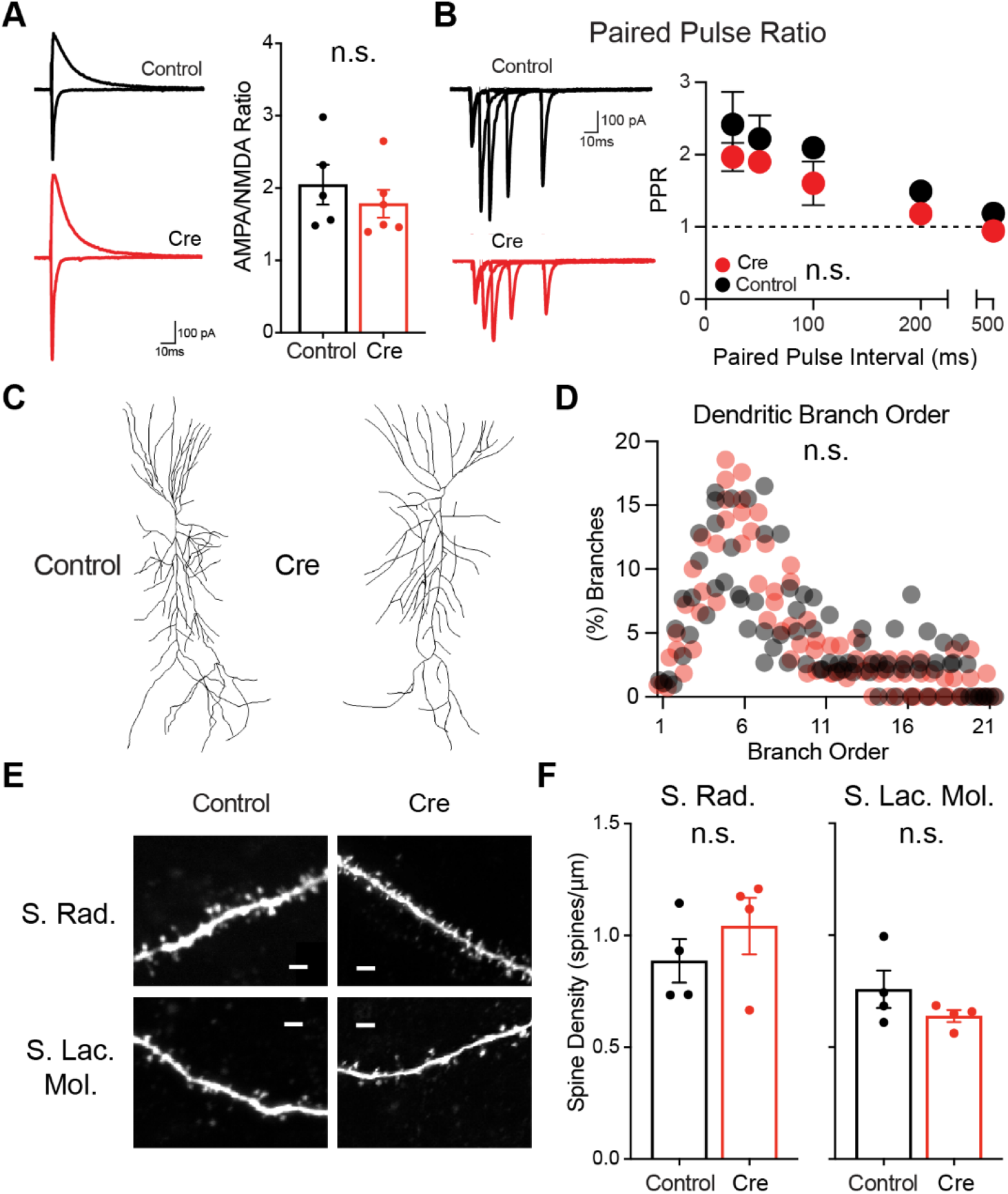
CA1-region *Stx3* deletion has no effect on basal synaptic transmission or dendritic properties. **A)** Measurements of postsynaptic AMPA and NMDA currents demonstrate that CA1-region deletion of *Stx3* does not affect basal ratios of AMPA to NMDA receptors. Left - representative traces, right - summary plot. [N=5 control cells, 6 Cre cells. Unpaired t-test: p=0.441] **B)** CA1-region deletion of *Stx3* does not affect presynaptic release probability as determined by the Paired Pulse Ratio (PPR). Left - representative traces, Right - summary plot. [N=5 control cells, 7 Cre cells. rmANOVA: inter-stimulus interval main effect p=0.0003, virus main effect p=0.061, interaction 0.968] **C)** Representative reconstruction of CA1-region neurons from biocytin filled cells. **D)** CA1-region deletion of *Stx3* does not affect gross dendritic morphology as demonstrated by the frequency distribution of branch order. [N=4 control cells, 4 Cre cells. The data are lognormally distributed, so we took the natural log and then performed mixed effects analysis. Branch order main effect p<10^−5^, virus main effect p=0.306] **E)** Representative 60x confocal dendritic spine images of biocytin filled CA-1 region neurons, scale bar: 2 µm. S. Rad = stratum radiatum, S. Lac. Mol = stratum lacunosum-moleculare. **F)** CA1-region deletion of Stx3 does not affect spine density in stratum radiatum (left) or stratum lacunosum-moleculare (right) as demonstrated by the average spine density measurements. [Unpaired t-test: N=4 control cells/40 dendritic segments, 4 Cre cells/40 dendritic segments for both sub-regions. Cells were used as the biological replicate. S. Rad. p=0.368, S. Lac. Mol p=0.221]

**Fig. S2.**
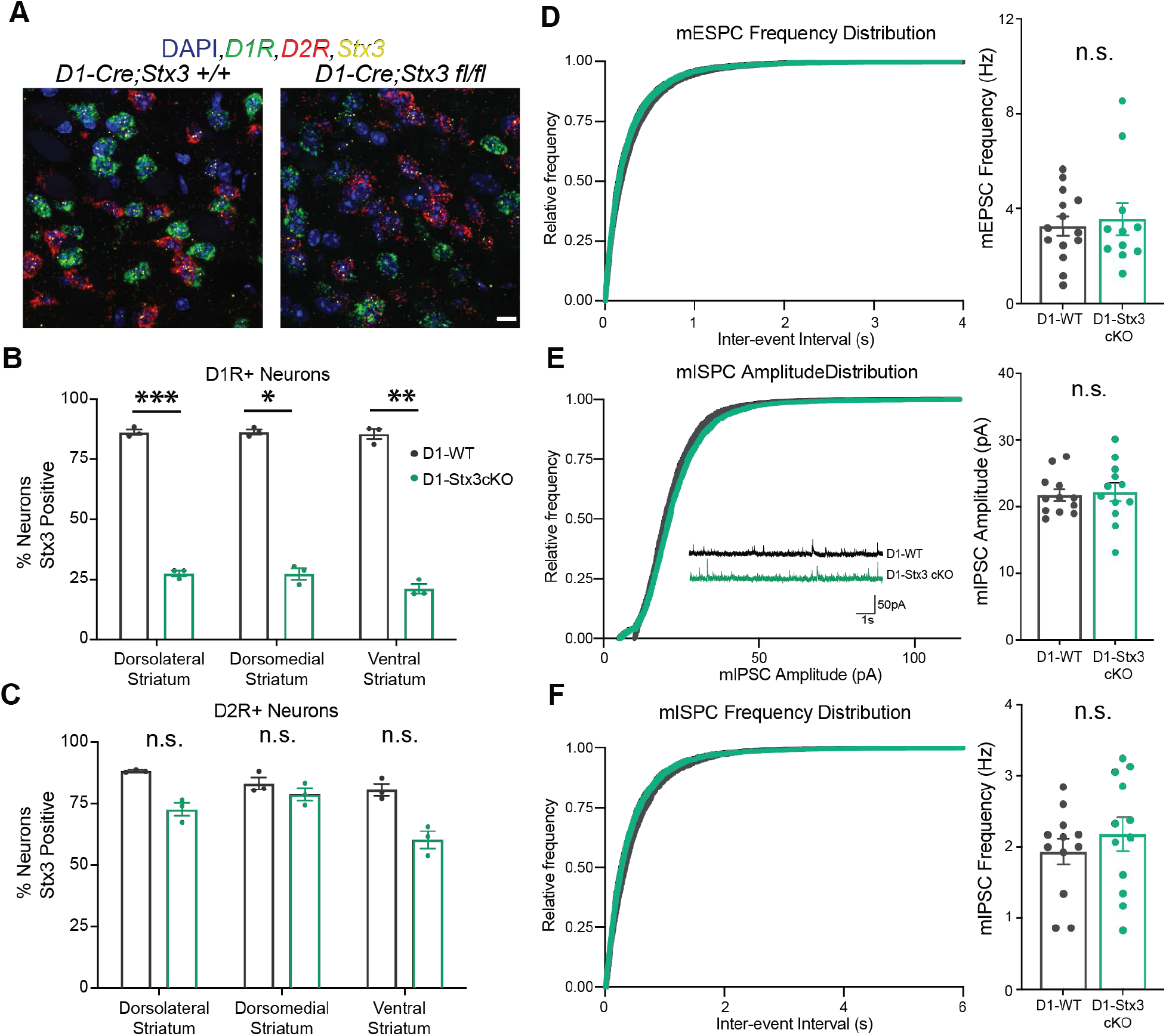
Characterization of D1-*Stx3cKO* neurons. **A)** Representative *in situ* hybridization images. Confocal images of coronal sections containing either Dorsolateral, Dorsomedial, or Ventral Striatum were analyzed for co-localization of *D1R, D2R,* and *Stx3* mRNA within the DAPI mask. Scale bar: 10 µm **B)** Summary plot of D1-*Stx3* cKO efficiency in D1R^+^ neurons. We observed a large decrease in D1R^+^/Stx3^+^ co-labeled neurons in the D1-*Stx3* cKO group. In order to account for multiple levels of nested repeated measures per biological replicate (i.e. different brain regions and cell-types from the same mouse) we used a 3-way ANOVA. [N=3 D1-WT mice, 3 D1-*Stx3cKO* mice (i.e. biological replicates). 18-32 sections per brain region (i.e. technical replicates), but biological replicates used for analyses. 3way ANOVA: brain region main effect p=0.0047, cell type main effect p<10^−5^, genotype main effect p<10^−5^, cell type x genotype interaction, p<10^−5^. The relevant adjusted p-values from multiple comparisons for *Stx3^+/+^* vs *Stx3^fl/fl^* within D1R^+^ neurons: Dorsolateral Striatum p=0.0002; Dorsomedial Striatum p=0.012; Ventral Striatum p=0.002] **C)** Same as (B), but for D2R+ neurons. These data demonstrate the cell-type specificity of the D1-*Stx3* cKO line. Continuation of multiple comparisons from the 3-way ANOVA for *Stx3^+/+^* vs *Stx3^fl/fl^* within D2R+ neurons: [Dorsolateral Striatum p=0.548; Dorsomedial Striatum p=0.973; Ventral Striatum p=0.345] **D)** D1-*Stx3* cKO does not affect basal excitatory synaptic frequency in striatal D1R^+^ neurons as demonstrated by cumulative frequency of miniature excitatory postsynaptic currents (mEPSC) inter-event-intervals (left) and summary plot of mEPSC frequencies (right). [N=14 D1-WT cells, 11 D1-*Stx3cKO* cells. Mann-Whitney: p=0.893] **E)** D1-*Stx3* cKO does not affect basal inhibitory synaptic strength in striatal D1R^+^ neurons as demonstrated by cumulative frequency of miniature inhibitory postsynaptic currents (mIPSC) amplitude (left, inset: representative mISPC traces) and summary plot of mIPSC amplitudes (right). [N=12 D1-WT cells, 12 D1-*Stx3cKO* cells. Unpaired t-test: p=0.787] **F)** same as (D) for mIPSCs. [N=12 D1-WT cells, 12 D1-*Stx3cKO* cells. Unpaired t-test: p=0.392] * indicates p<0.05, ** indicates p<0.01, *** indicates p<0.001

**Fig. S3.**
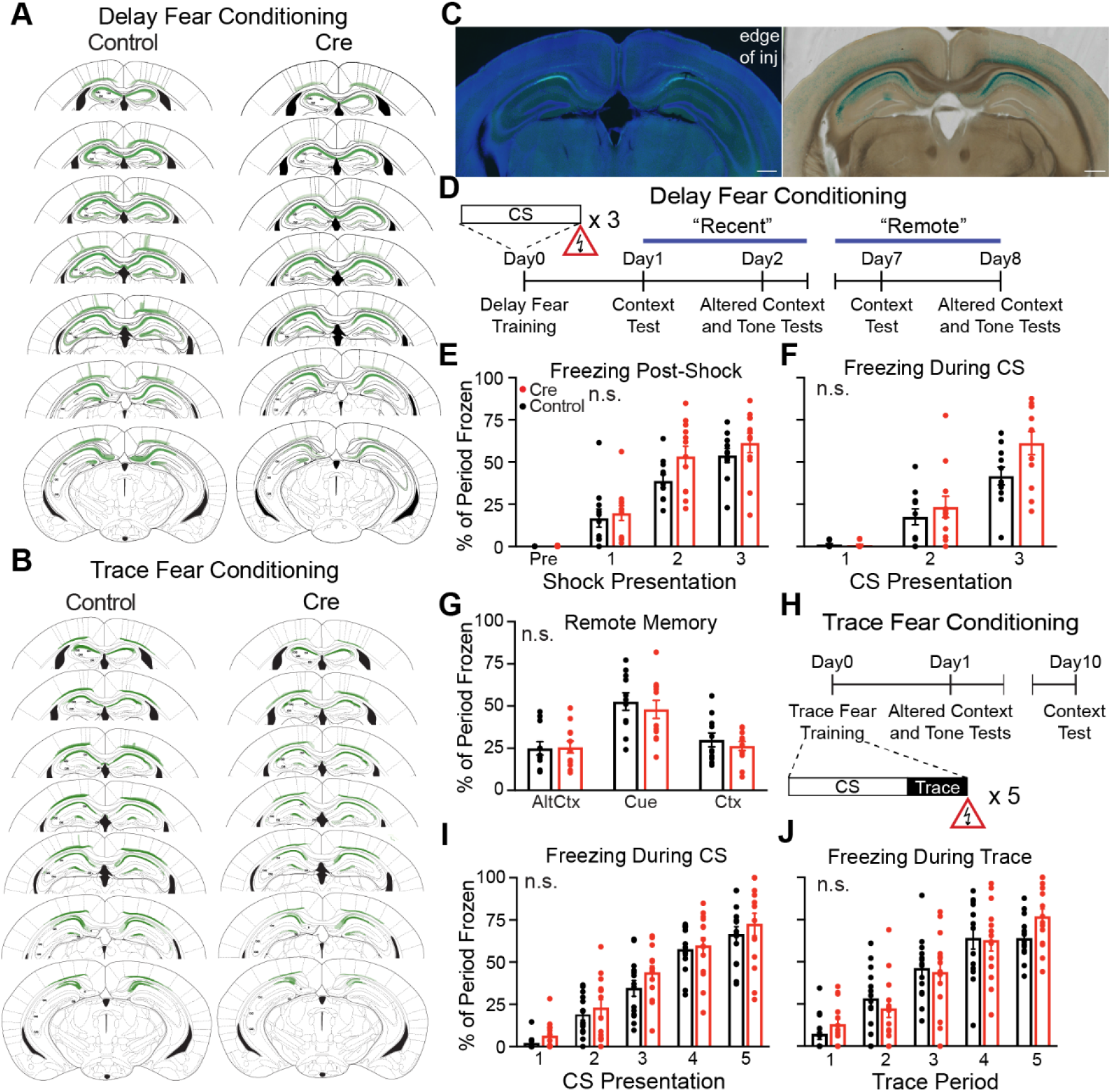
Expanded Fear Conditioning data. **A)** Summary plot of viral spread for every mouse that underwent Delay Fear Conditioning. Each mouse is drawn with a semi-transparent fill – darker green indicates more overlap across mice. **B)** Same as (A) for Trace Fear Conditioning **C)** Quantification of viral spread based on EGFP fluorescence underestimates extent of Cre recombination. Consecutive 100 µm thick sections of a Cre-injected *Stx3^fl/fl^* mouse at the posterior edge of EGFP expression. Left - EGFP (green) with DAPI counterstain (blue), Right - Ꞵ-galactosidase chromogenic stain. Ꞵ-galactosidase is only expressed upon Cre recombination. Scale bars: 500 µm **D)** Schematic of the Delay Fear Conditioning procedure, applies to panels E-G. **E)** Percent of the period spent frozen during the baseline and 1 min after each shock. This serves as a proxy for shock sensitivity. [N=11 control mice, 12 Cre mice. rmANOVA: Shock Number main effect p<10^−5^, virus main effect p=0.153, interaction p=0.227] **F)** Percent of each tone spent frozen during training demonstrates that both groups encoded the cue-shock association. [N=11 control mice, 12 Cre mice. rmANOVA: Cue Number main effect p<10^−5^, virus main effect p=0.073, interaction p=0.098] **G)** “Remote memory” tests for context, altered context, and cue-induced freezing. Testing occurred 1 week after conditioning. All mice previously experienced the “Recent” memory test, and essentially went through extinction training during those tests. [N=11 control mice, 12 Cre mice. rmANOVA: Test Condition main effect p<10^−5^, virus main effect p=0.557, interaction p=0.753] **H)** Schematic of the Trace Fear Conditioning procedure, applies to panels I & J. **I)** Percent of each tone spent frozen during training demonstrates that both groups encoded the cue-shock association. [N=14 control mice, 14 Cre mice. rmANOVA: Cue Number main effect p<10^−5^, virus main effect p=0.181, interaction p=0.916] **J)** Percent of each 15s trace period spent frozen. [N=14 control mice, 14 Cre mice. rmANOVA: Trace Number main effect p<10^−5^, virus main effect p=0.632, interaction p=0.28]

**Fig. S4.**
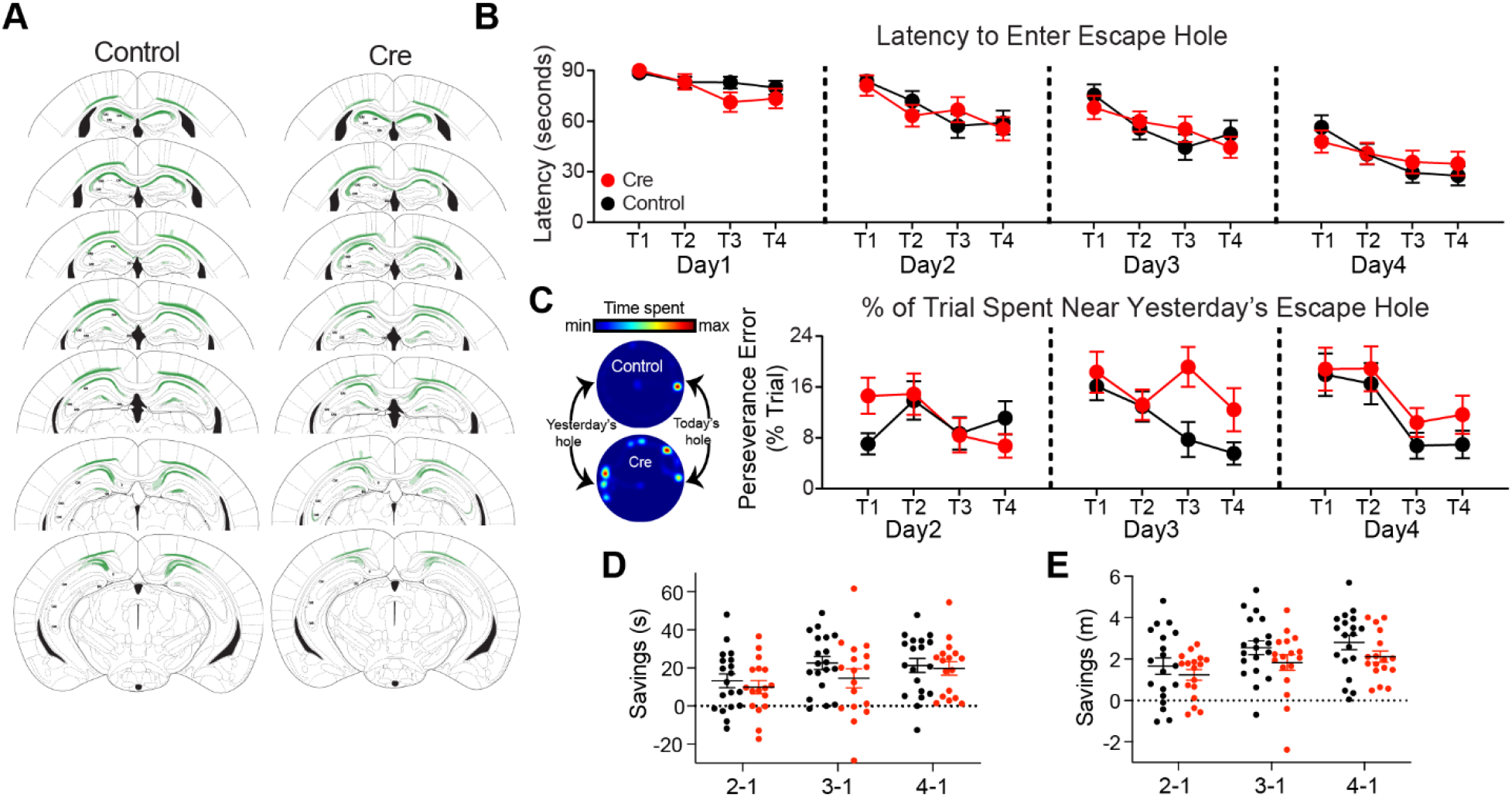
Expanded DMP Barnes data. **A)** Summary plot of viral spread for every mouse that underwent DMP Barnes. Each mouse is drawn with a semi-transparent fill – darker green indicates more overlap across mice. **B)** Latency to enter the proper escape port demonstrates that CA1-region LTP is not required for spatial learning, and daily re-learning. [N=19 control mice, 17 Cre mice. rmANOVA: trial main effect p<10^−5^, virus main effect p=0.779, interaction p=0.663] **C)** Example occupancy plot of a mouse on day 3, trial 3 (left) and % of trial on days 2-4 spent near yesterday’s escape hole (right, i.e. perseverance). While there is a main effect of virus, none of the post-hoc comparisons between genotypes on individual trials are significant. Further, this difference in perseverative errors did not affect the overall latency, path length, or savings - suggesting that CA1-region LTP does not affect overall performance. [N=19 control mice, 17 Cre mice. rmANOVA: trial main effect p=0.0003, virus main effect p=0.005, interaction p=0.32. Post-hoc comparison control vs Cre with Holm-Šídák correction all not significant] **D)** Savings of the latency to enter the escape port shows no difference between groups - indicating similar learning. “2-1” is calculated by subtracting latency on trial 2 from latency on trial 1 and averaging across all days. This can be interpreted as the amount of improvement or learning from trial 1 to 2. This extends to 3-1 and 4-1. [N=19 control mice, 17 Cre mice. rmANOVA: interval main effect p=0.005, virus main effect p=0.335, interaction p=0.474] **E)** Same as (D) but for savings of path length. [N=19 control mice, 17 Cre mice. rmANOVA: interval main effect p<10^−5^, virus main effect p=0.143, interaction p=0.763] * indicates p<0.05

**Fig. S5.**
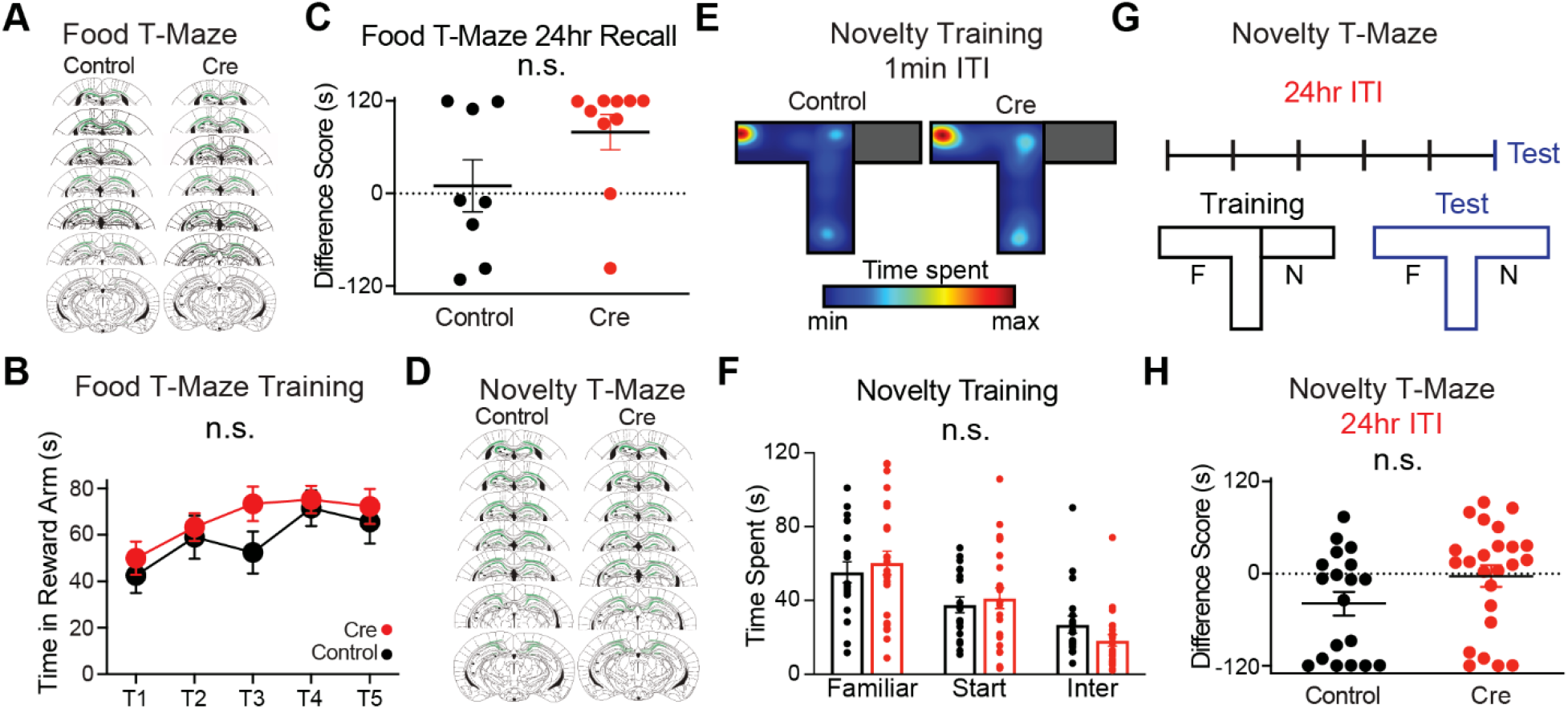
Expanded T-Maze data. **A)** Summary plot of viral spread for every mouse that underwent Food T-Maze **B)** CA1-region deletion of *Stx3* does not affect food T-Maze training, as demonstrated by the summary plot of time in the reward arm for each trial. [N=15 control mice, 21 Cre mice. mixed effects ANOVA: trial main effect p=0.0005, virus main effect p=0.29, interaction p=0.568] **C)** Summary plot of the difference score in the Food T-Maze 24hrs after the final test. Values were averaged across 2 test sessions, 1 min apart. [Unpaired t-test: N=8 control mice, 10 Cre mice. unpaired t-test p=0.096] **D)** Summary plot of viral spread for every mouse that underwent Novelty T-Maze **E)** Occupancy plot of representative mice during novelty T-Maze training. **F)** CA1-region deletion of *Stx3* does not affect novelty T-Maze training, as demonstrated by the summary plot of time spent in training T-Maze compartments (averaged over all trials for each mouse). Although there was a large effect during the 1min ITI test, both groups of mice occupied the 3 zones of the maze during training at similar rates. “Inter” = intermediate zone, the zone at the intersection of all arms. [N=19 control mice, 24 Cre mice. rmANOVA: compartment main effect p<10^−5^, virus main effect p=0.81, interaction p=0.51] **G)** Schematic of the 24hr T-Maze procedure. The training configuration is depicted in black – with a separator blocking one arm (side counterbalanced across groups). 5 training trials with a 24hr ITI were followed by a 24hr ITI and test trial (depicted in blue, no separators). **H)** Both groups performed at chance levels with a 24hr ITI - summary plot. Difference Score = Time spent in (Novel)-(Familiar). Dotted line indicates equivalent/chance-level exploration (0s). [N=19 control mice, 24 Cre mice. Mann-Whitney: p=0.067]

**Fig. S6.**
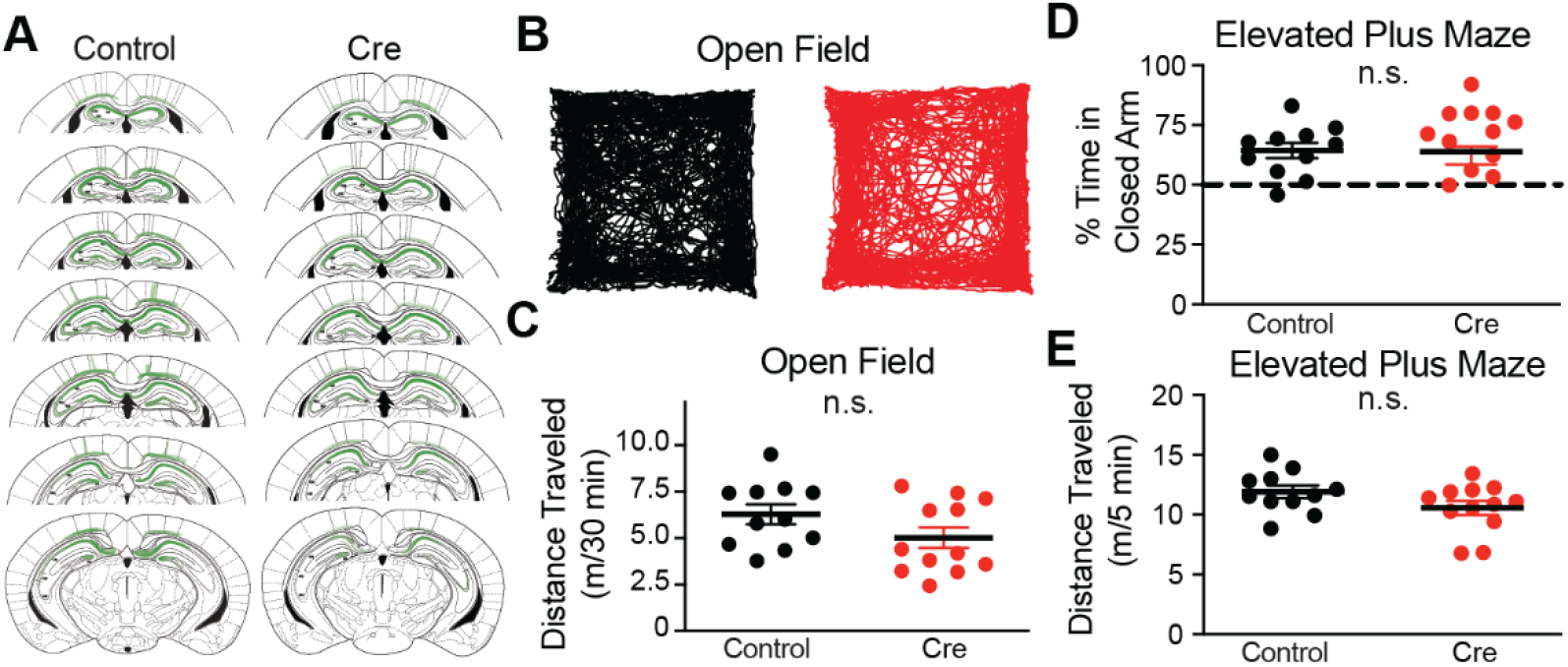
CA1-region *Stx3* is not involved in basal locomotion and anxiety-like behavior. **A)** Summary plot of viral spread for every mouse that underwent Open Field and Elevated Plus Maze. **B)** Representative open field tracks. **C)** CA1-region deletion of *Stx3* does not affect basal locomotion as demonstrated by the summary plot of distance traveled in the open field. [N=11 control mice, 12 Cre mice. Unpaired t-test: p=0.116] **D)** CA1-region deletion of *Stx3* does not affect anxiety-like behavior as demonstrated by the summary plot of %time spent in the closed arm of the elevated plus maze. [N=11 control mice, 12 Cre mice. Unpaired t-test: p=0.924] **E)** CA1-region deletion of *Stx3* does not affect locomotion in the elevated plus maze as demonstrated by the summary plot of distance traveled. [N=11 control mice, 12 Cre mice. Unpaired t-test: p=0.107]

**Fig. S7.**
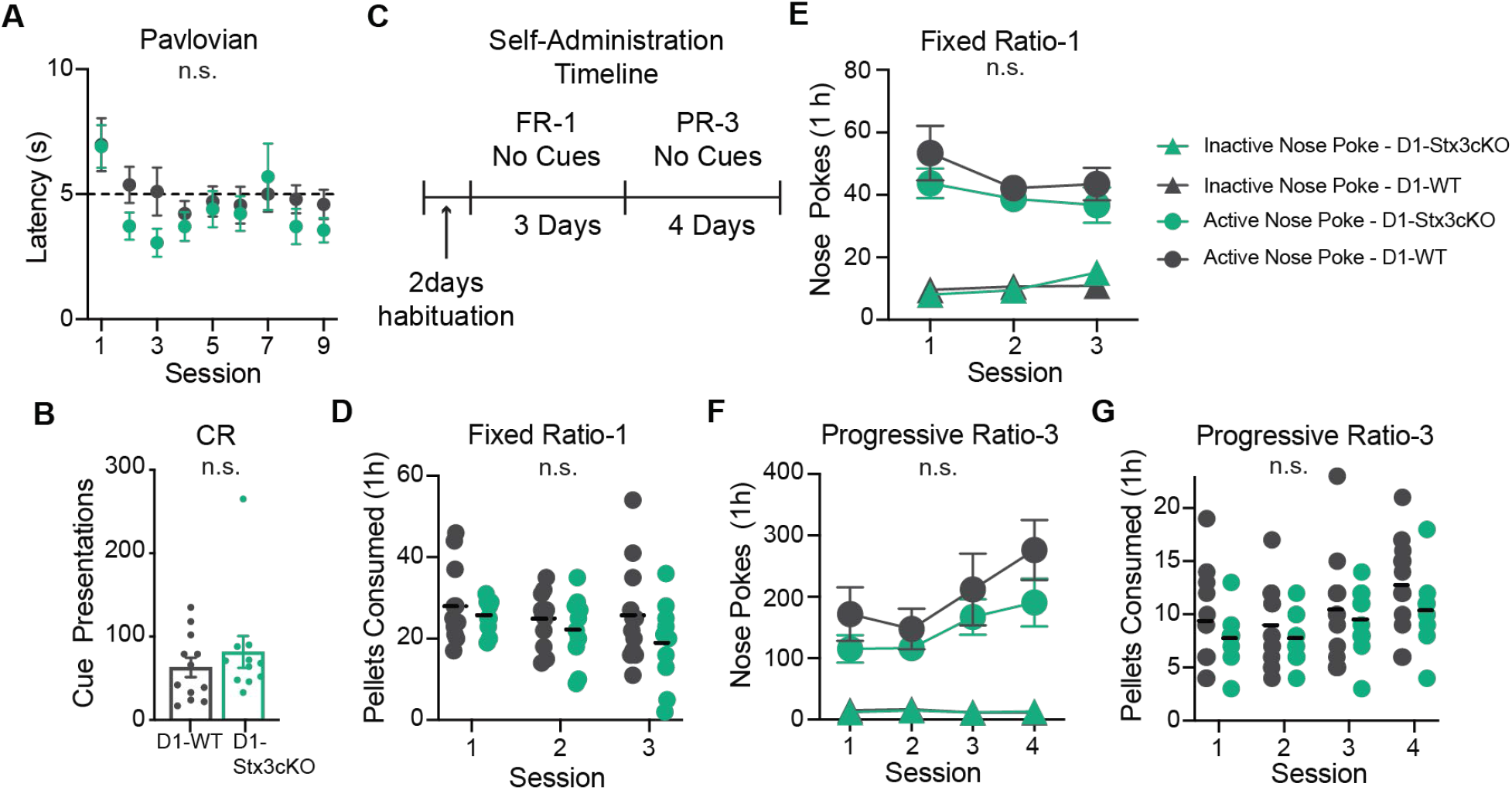
Expanded Pavlovian and Self-Administration Data. **A)** LTP in D1R^+^ neurons does not affect learning the timing of reward delivery as demonstrated by latency of port entry (PE) after cue onset. Horizontal dashed line at 5 s represents reward delivery. [N=12 D1-WT mice, 11 D1-*Stx3cKO* mice. rmANOVA: session main effect p=0.003, genotype main effect p=0.302, interaction p=0.506] **B)** LTP in D1R^+^ neurons is not involved in learning to nose-poke for presentations of the conditioned stimulus (CS) used in Pavlovian conditioning. The number of cues (conditioned reinforcers, CR) acquired during the conditioned reinforcement test is plotted. [N=12 D1-WT mice, 11 D1-*Stx3cKO* mice. Mann-Whitney: p=0.458] **C)** Food self-administration schematic - all testing occurred after Pavlovian conditioning and the conditioned reinforcement (CR) test. Mice received 1 day of “pre-training” with cues and food restriction, and another day of “pre-training” with cues but without food restriction. Then cues were removed for the remaining experiments to study the hedonic properties of the food itself. Note these mice were fed *ad libitum* in the homecage. Each operant chamber contained an “active” and “inactive” nose-poke port. Nose-pokes into the active port led to food delivery (1 nose-poke required per pellet i.e. fixed ratio-1) while nose-pokes into the inactive port had no consequences. During progressive ratio-3 testing, the response requirement for one food pellet began at 1 nose-poke, and increased by 3 nose-pokes for each subsequent food pellet (i.e. 1+4+7 nose-pokes required to receive 3 food pellets). **D)** Fixed ratio-1 training demonstrated that LTP in D1R^+^ neurons is not involved in stable food self-administration under a low response requirement. Pellets consumed each session are plotted here. Pellets earned but not consumed were not counted in this measure. [N=12 D1-WT mice, 10 D1-*Stx3cKO* mice. rmANOVA session main effect p=0.07, genotype main effect p=0.226, interaction p=0.421] **E)** LTP in D1R^+^ neurons is dispensable for learning to discriminate between nose-poke ports as demonstrated by the summary plot of active and inactive nose-pokes during fixed ratio-1 training. [N=12 D1-WT mice, 10 D1-*Stx3cKO* mice. Three-way ANOVA matching for session and port: session main effect p=0.43, port main effect p<10^−5^, genotype main effect p=0.407, session x port interaction p=0.039, all other interactions n.s. The only post-hoc tests with Holm-Šídák adjusted p-value below threshold were comparisons between active and inactive nose-poke ports] **F)** The progressive ratio-3 test revealed that LTP in D1R^+^ neurons is dispensable for adapting response requirements under increasing demands. The summary plot of active and inactive nose-pokes during progressive ratio-3 training is shown here. These data again demonstrate the ability of both groups to discriminate between the active and inactive nose-poke ports. These data can be averaged over sessions to produce Fig 2M. [N=12 D1-WT mice, 10 D1-*Stx3cKO* mice. Three-way ANOVA matching for session and port: session main effect p<10^−5^, port main effect p<10^−5^, genotype main effect p=0.321, session x port interaction p=0.0001, all other interactions n.s. The only post-hoc tests with Holm-Šídák adjusted p-value below threshold were comparisons between active and inactive nose-poke ports] **G)** LTP in D1R^+^ neurons does not play a role in food self-administration under a progressive ratio-3 response requirement as demonstrated by the summary plot of pellets consumed over sessions. [N=12 D1-WT mice, 10 D1-*Stx3cKO* mice. rmANOVA session main effect p<10^−5^, genotype main effect p=0.313, interaction p=0.697]

**Fig S8.**
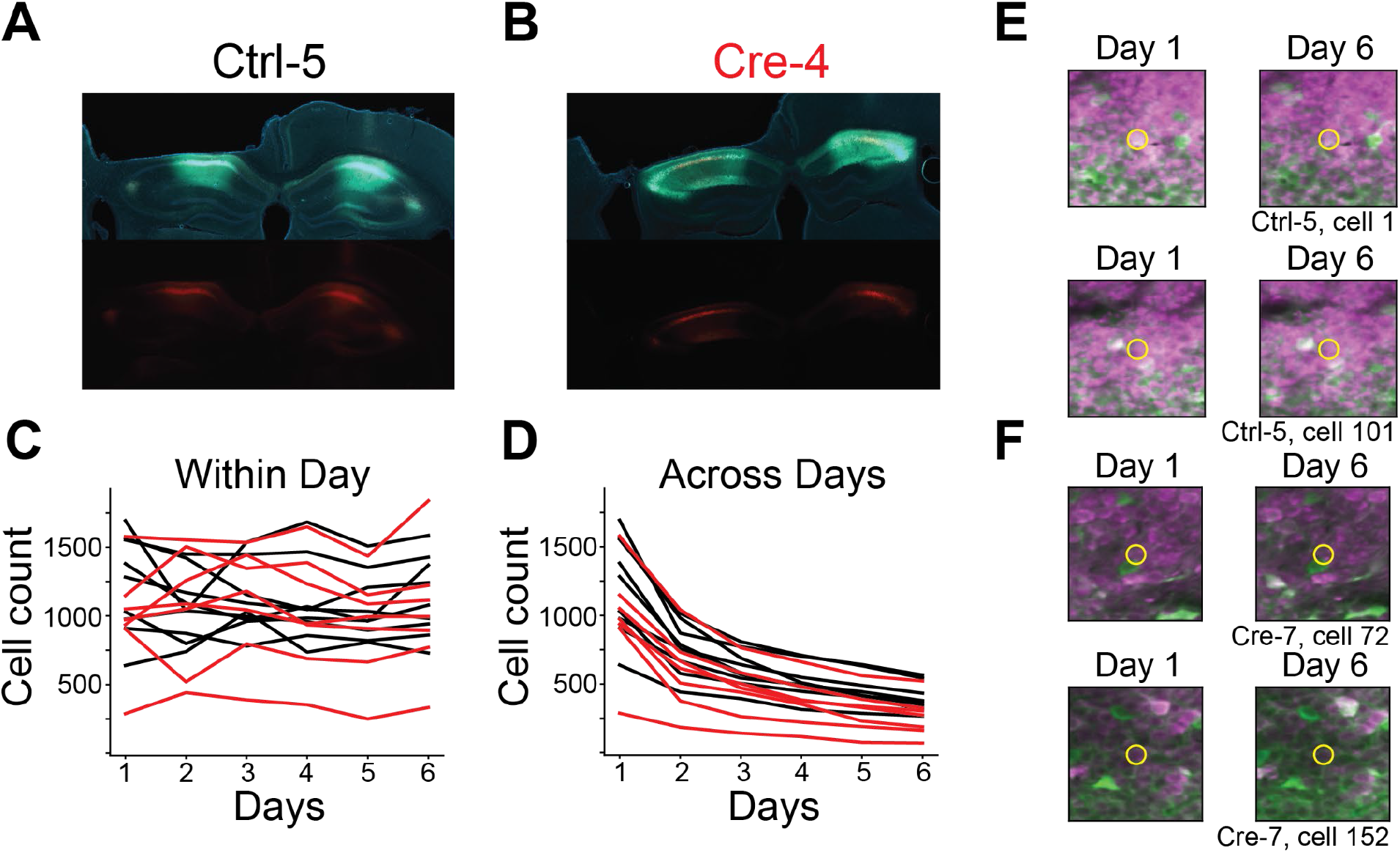
Virus coexpression and two photon cell tracking. **A)** Example histology from an example control mouse (Ctrl-5). Coronal slice showing unilateral imaging implant and bilateral virus expression. Top - Coexpression of DAPI (blue), GCaMP (green), and mCherry (red). Bottom - Expression of mCherry. **B)** Same as (A) for an example Cre mouse (Cre-4). **C)** Number of cells recorded in each session for each mouse (black - control, red - Cre). Only GCaMP^+^/mCherry^+^ neurons were included in all analyses. **D)** Number of cells tracked across all sessions using ROIs from day 1. **E)** Spatial footprints of example cells tracked over 6 days of imaging from Fig 3E. 2-photon mean image from day 1 (left) and day 6 (right, magenta-mCherry, green-GCaMP). Yellow circle highlights the tracked cell. **F)** Same as (E) for cells from Fig 3G

**Fig. S9.**
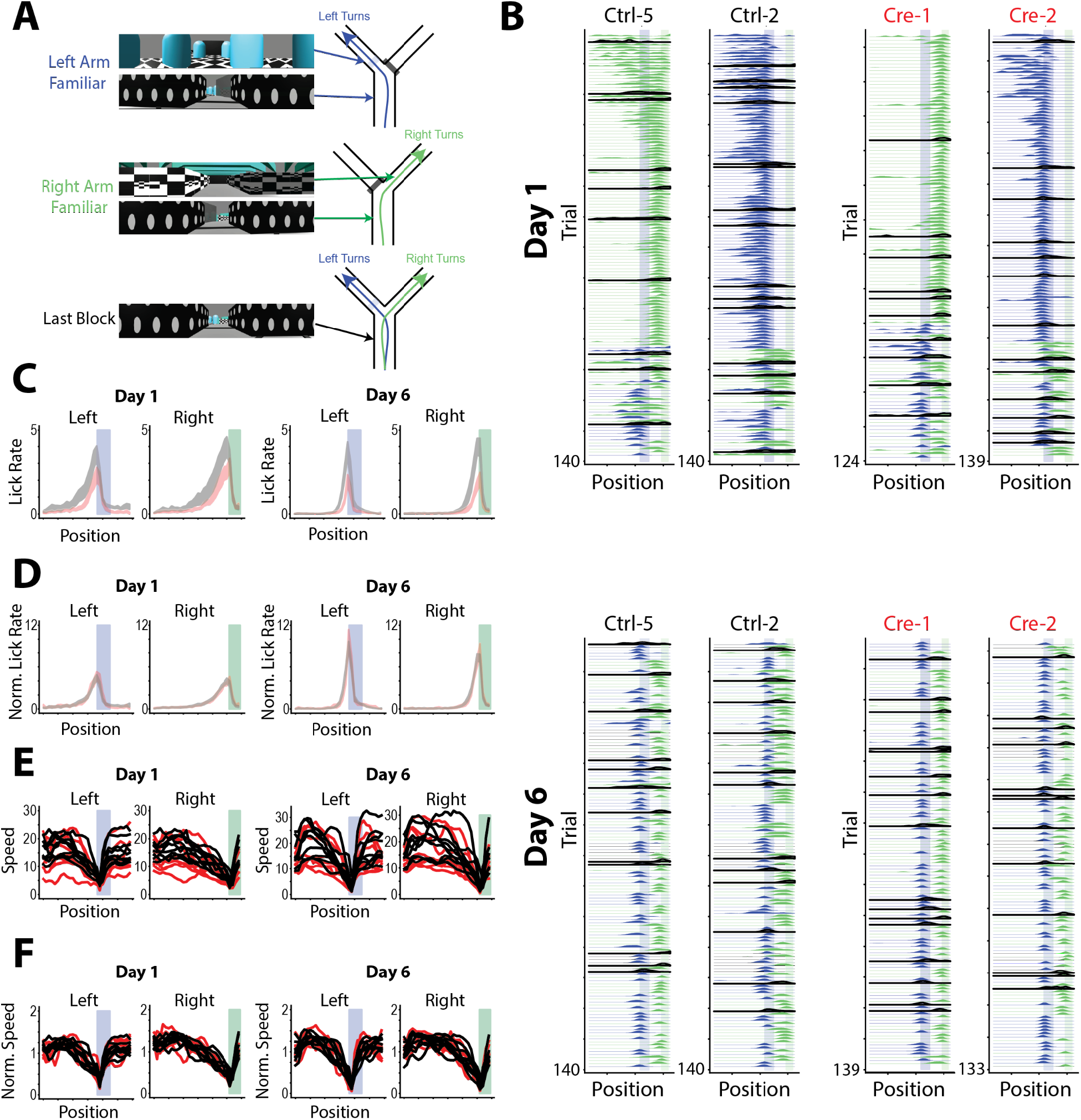
Expanded VR description and behavior. **A)** Example screenshots of virtual reality track for left arm familiar trials (top), right arm familiar trials (middle) and the stem of trials from the last block where the virtual curtain is removed. We counterbalanced whether the right or left arm was the familiar arm across animals. **B)** Smoothed lick rate on each trial as a function of position for example mice (Top Row - Day 1 examples, Bottom Row - Day 6 examples). Trials are colored by whether they are a left trial (blue) or right trial (green). Black highlighted trials are probe trials in which the reward was omitted. Reward zones are shown by the vertical shaded regions (blue - left reward zone, green - right reward zone). Right arm familiar (Ctrl-5 and Cre-1) and left arm familiar (Ctrl-2 and Cre-2) mice are shown. **C)** Mean lick rate as a function of position for left and right trials for days 1 and 6. Reward zones are highlighted as in (B). This serves as a control analysis to ensure across group differences in lick rates are not driven by differences across the virtual environments. Unlike Fig 3C-D, both novel and familiar trials are intermixed here (black - control, red - Cre, shaded regions indicate across animal mean ± SEM). **D)** Mean normalized lick rate as a function of position for left and right trials for days 1 and 6. Each mouse’s position-binned lick rate is divided by the mouse’s overall average lick rate. Reward zones are highlighted as in (B) (black - control, red - Cre, shaded regions indicate across animal mean ± SEM) **E)** Mean running speed as a function of position for left and right trials for days 1 and 6 (black - control, red - Cre, each line is different mouse) **F)** Mean normalized running speed as a function of position for left and right trials on days 1 and 6. Each mouse’s position-binned running speed is divided by the mouse’s overall average running speed. The characteristic decrease in running speed to consume the reward can be seen most clearly here. (black - control, red - Cre, each line is a different mouse)

**Fig. S10.**
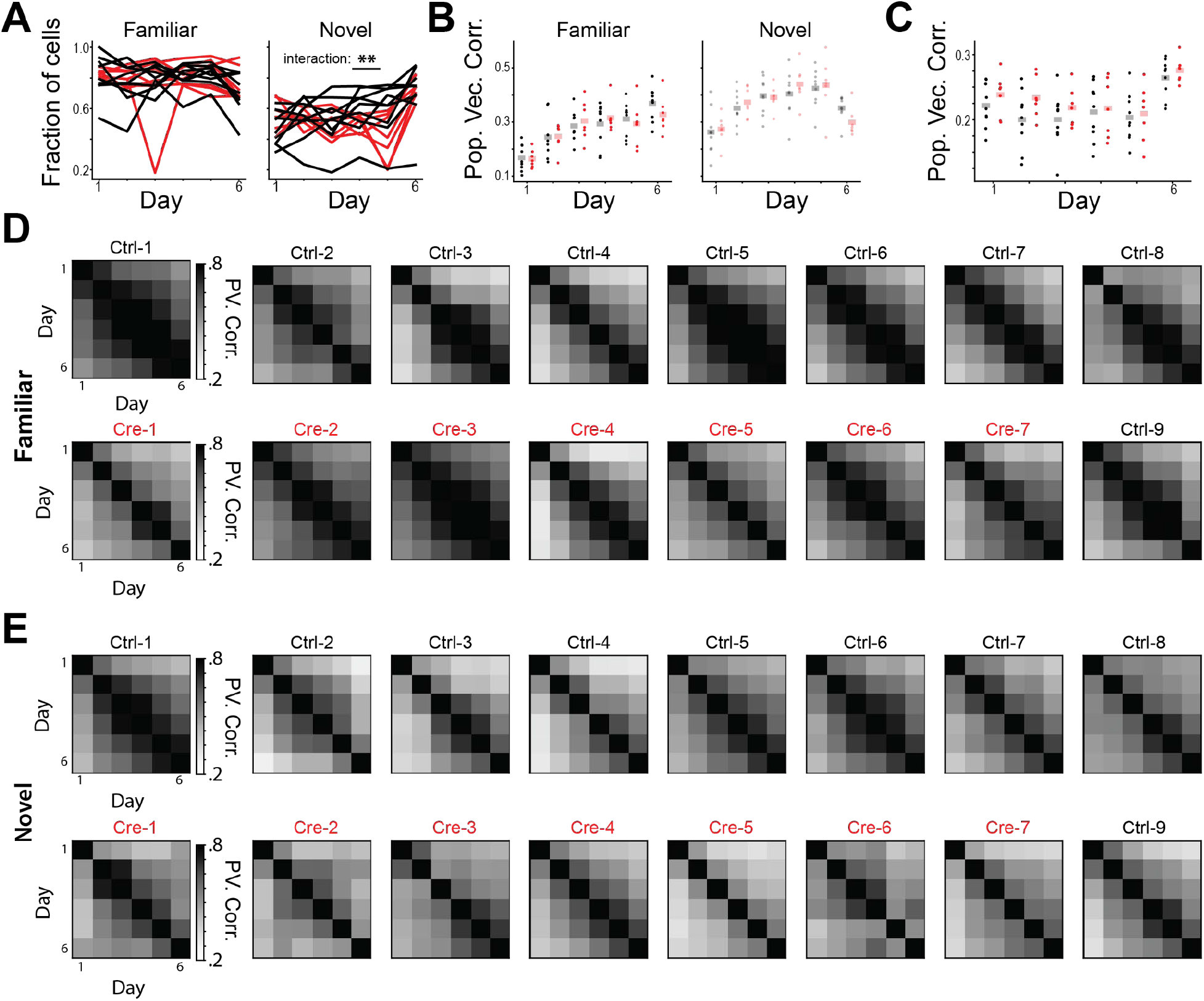
Place cell population stability and remapping. **A)** Fraction of cells with significant spatial information on each day for familiar (left) and novel (right) trials. Each line represents data from a single animal. We found a significant interaction between virus and day for novel trials but none of the posthoc tests were significant. [N=9 control mice, 7 Cre mice, 6 days. Familiar trials mixed effects ANOVA: virus main effect p=0.999, day main effect p=0.459, interaction p=0.605. Novel trials mixed effects ANOVA: virus main effect p=0.487, day main effect p=0.516, interaction p=0.002, all posthoc tests p>0.05]. **B)** Average trial x trial population vector correlation for familiar (left) and novel (right) trials on each day shows similar within-day stability between conditions. Each dot is the average for a single animal. [N=9 control, 7 Cre mice, 6 days. Familiar trials mixed effects ANOVA: virus main effect p=0.899, day main effect p=1.52×10^−17^, interaction p=0.342. Novel trials mixed effects ANOVA: virus main effect p=0.907, day main effect p=3.03×10^−20^, interaction p=0.025, all posthoc tests p>0.05] **C)** Population vector correlation between the average familiar trial activity and the average novel trial activity on each day shows comparable remapping across groups. [N=9 control, 7 Cre animals, 6 days. Mixed effects ANOVA: virus main effect p=0.300, day main effect p=1.06×10^−8^, interaction p=0.71] **D)** Day x day population vector correlation for familiar trials for all mice demonstrates comparable long term stability across groups. **E)** Same as (D) for novel trials.

**Fig. S11.**
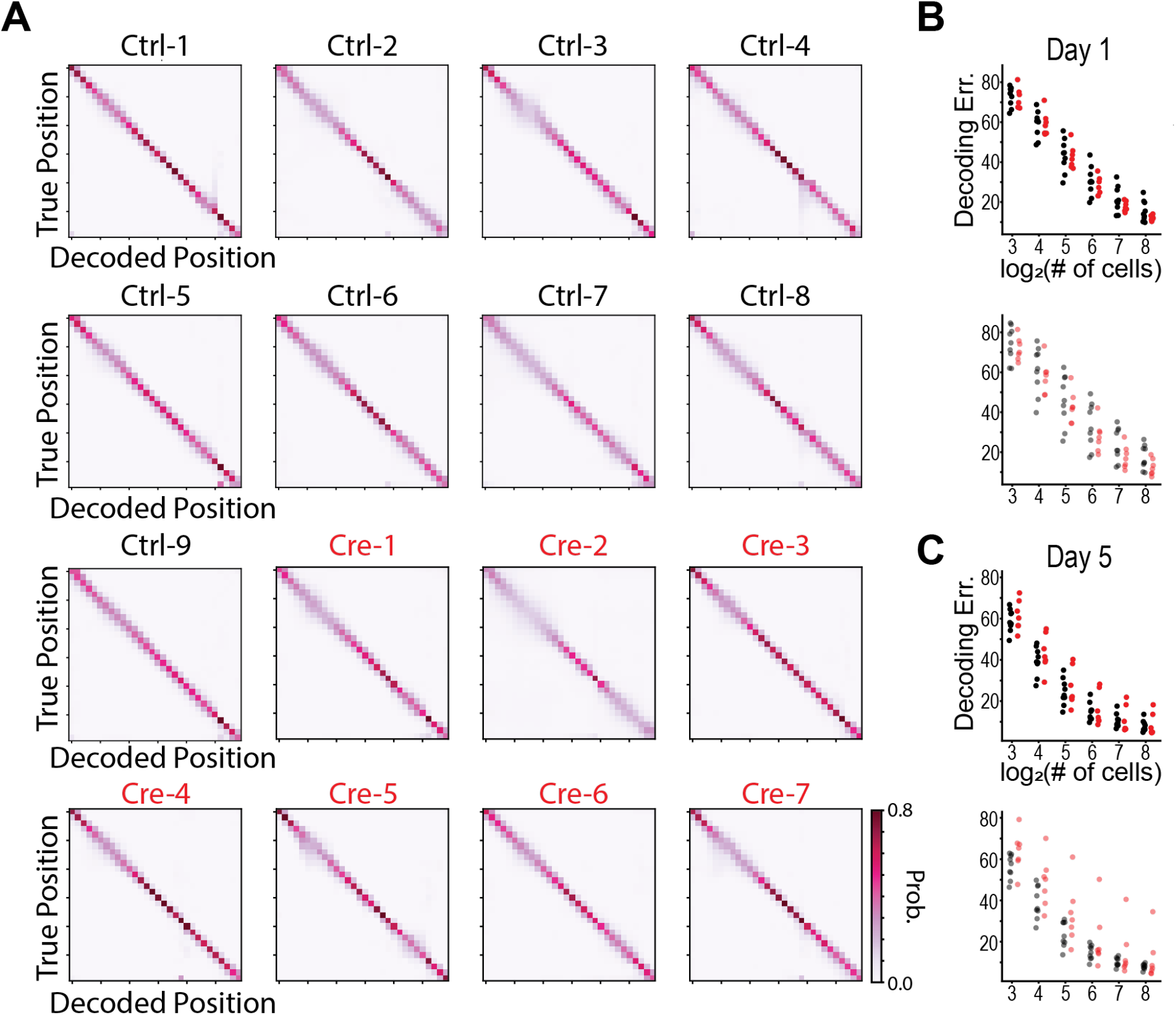
LTP is not required to form accurate and and decodable population spatial codes. **A)** Leave-one-trial-out average posterior probability for Naive Bayes position decoder trained and tested on day 1 familiar trial activity for each animal. The majority of probability mass lies along the diagonal indicating accurate decoding. **B)** Mean absolute deviation between the true position and decoded position from leave-one-trial-out cross-validation for Naive Bayes models trained with different numbers of cells on day 1 data. Each dot is the average error of 50 models trained with cells randomly chosen with replacement for a single animal (top - familiar trials, bottom - novel trials). **C)** Same as (B) for day 5 trials.

**Fig. S12.**
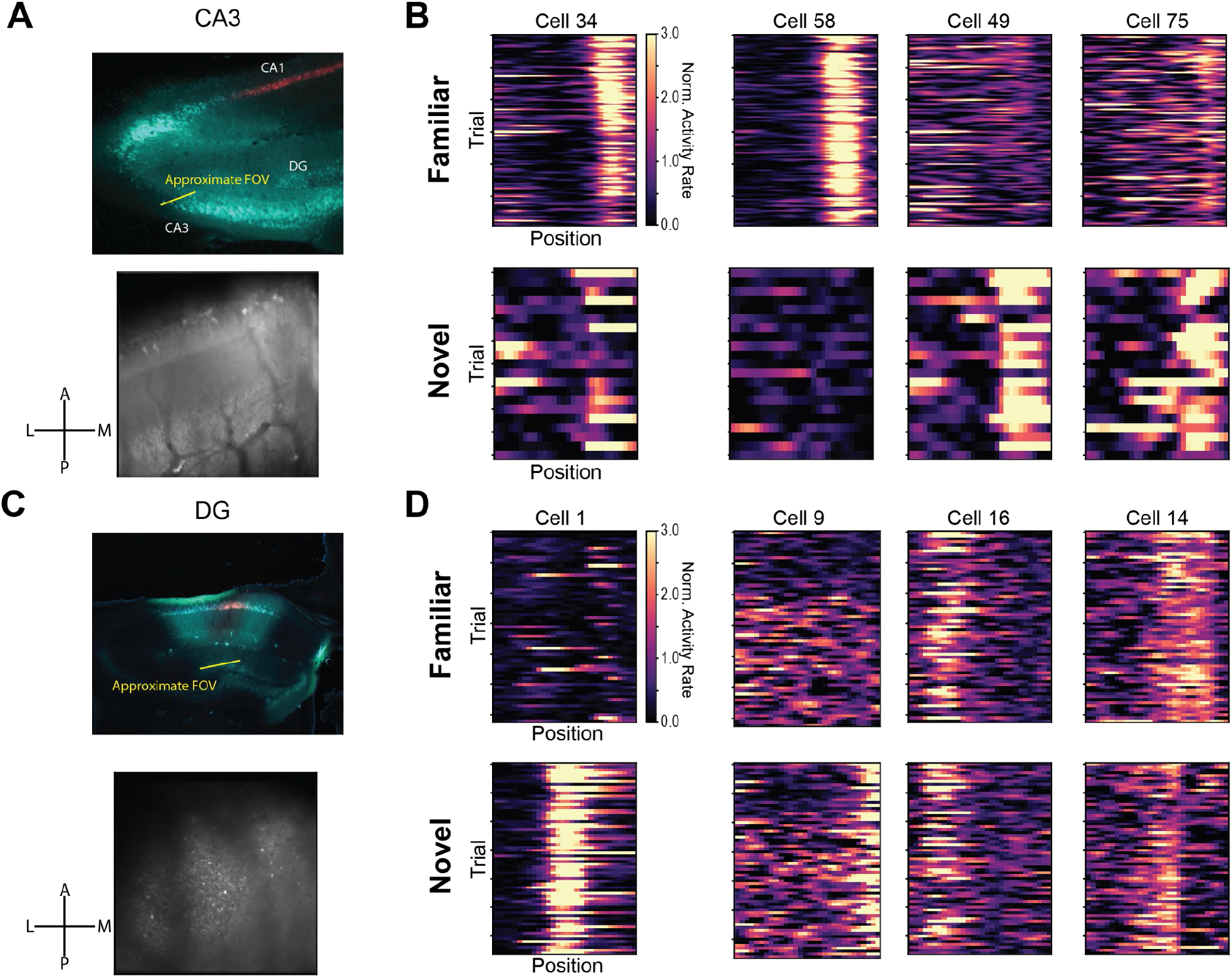
Intact place cells in CA3 and DG from Cre animals. **A)** Top *-* histology from a mouse in which we were able to image CA2/3 place cells (Cre-8,DAPI-blue, GCaMP- green, mCherry - red). Bottom *-* the average two photon field of view from an example imaging session. Note the visible ventricle on the anterolateral edge of the 2-photon FOV. Crosshairs denote anterior-posterior (A-P) and lateral-medial (L-M) axes in FOV **B)** Example trial x position activity rate maps from CA2/3 place cells on day 1 for familiar (top) and novel (bottom) trials. Each column of plots indicates a different cell. **C)** Top *-* histology from a mouse in which we were able to image dentate gyrus (DG) place cells (Cre-7, DAPI-blue, GCaMP-green, mCherry - red). Bottom *-* the average two photon field of view. We were able to image DG by focusing ∼600 microns past the CA1 imaging FOV. **D)** Example trial x position activity rate maps for DG place cells on an example session. This session was performed after all other training and was a repeat of the procedure from day 6 (Fig 3B).

**Fig. S13.**
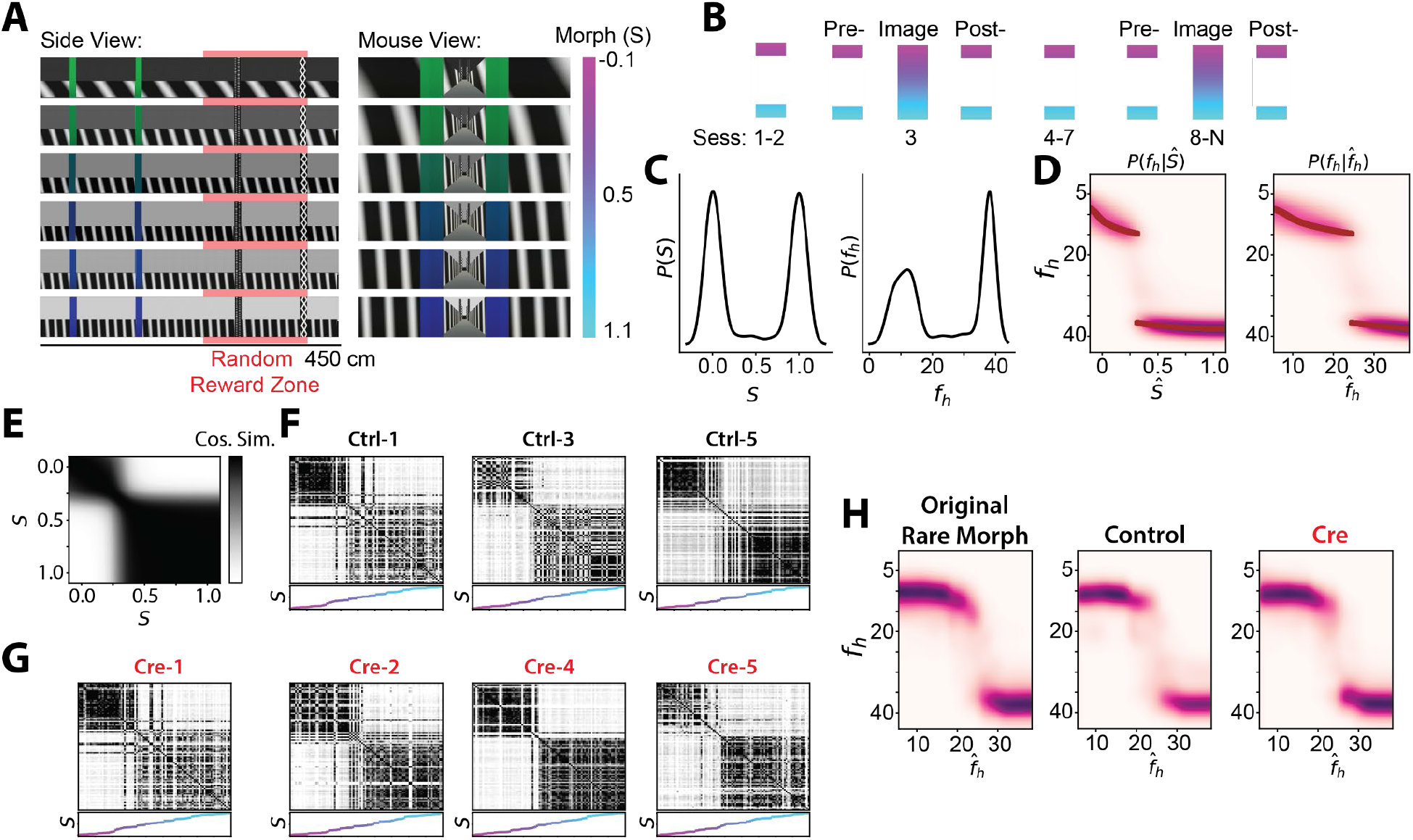
Experience dependent hippocampal remapping does not require CA1 LTP. **A)** See Supplemental Text “Experience dependent hippocampal remapping” for more details. Example VR tracks that are gradually “morphed” between two extremes (S-morphing parameter; side view - left, mouse view - right). Rewards were placed at random locations on the back half of the track on each trial (red highlighted region). **B)** Training protocol for inducing discrete remapping to intermediate morph values. Animals experienced randomly interleaved tracks of the extreme morph values except during a subset of imaging sessions (days 3 and 8-N). During these imaging sessions, morph values from the full range of stimuli were randomly interleaved. **C)** Left - Example prior distribution over morph values prior to session 8 for a mouse trained under the protocol in (B). Right - Same prior distribution after converting morph values to frequency of wall stimuli (*f_h_*). **D)** Posterior distributions, 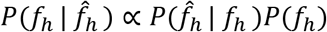, for each morph value (left) or each value of wall frequency (right) for an animal performing optimal Bayesian inference under the prior shown in (C). If the hippocampus remapped to reflect the animal’s best estimate of which environment it was occupying, remapping patterns should encode this distribution. **E)** Trial x trial cosine similarity matrix for a simulated population that encodes the posterior distribution exactly. **F)** Example trial x trial population cosine similarity matrices for session 8 for each control mouse **G)** Example trial x trial population cosine similarity matrices for session 8 for each Cre mouse. **H)** Left - Unsupervised reconstruction of posterior distribution from neural data in (*50*). Middle - reconstruction of posterior distribution from control mice. Right - Reconstruction of posterior distribution from Cre mice. Panels A-D and H (left) reproduced from (*50*).

**Fig. S14.**
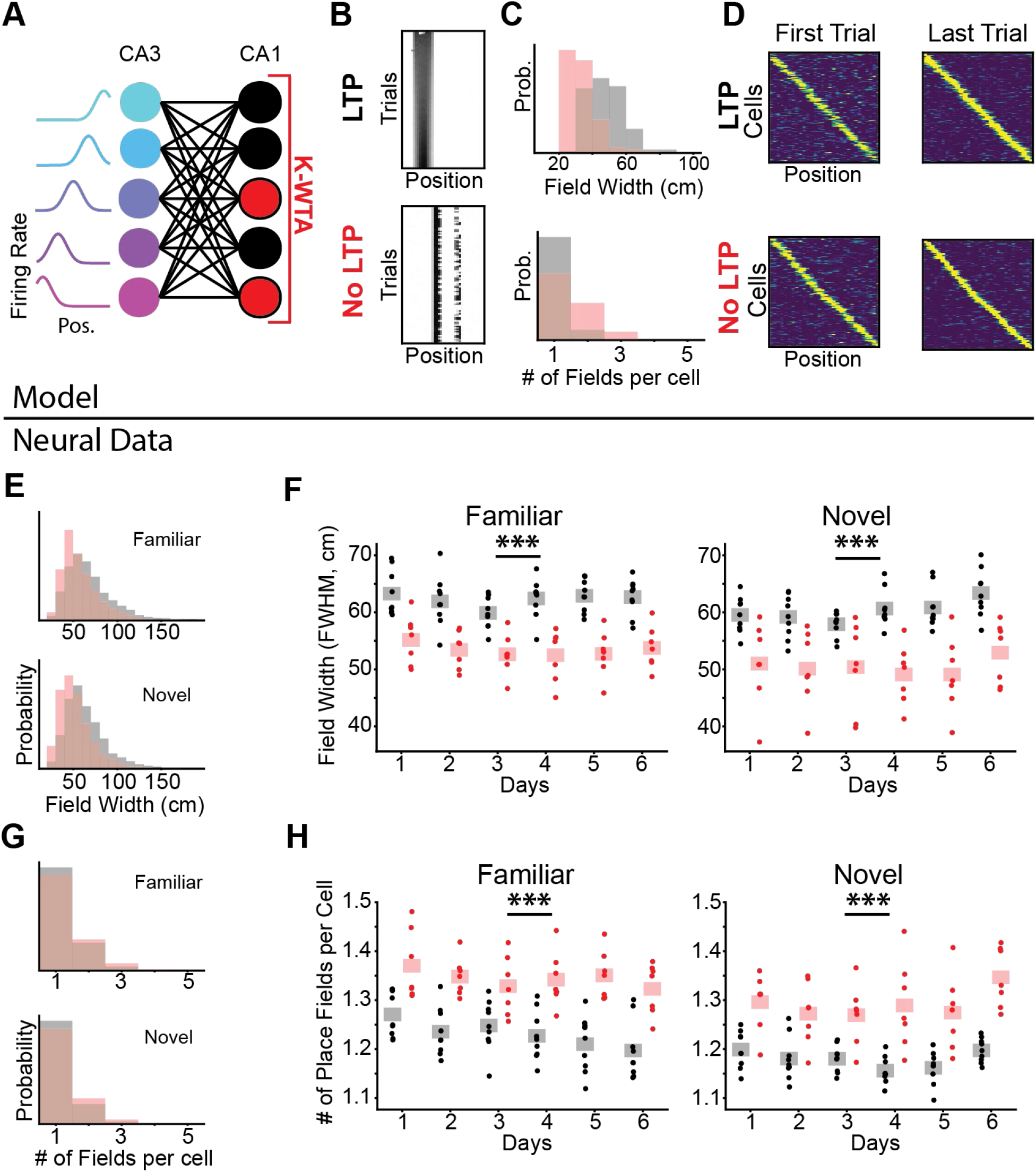
Model for passively inherited versus learned place cell representations explains differences in place cell properties between control and Cre mice. **A)** Schematic for K-Winners-Take-All (KWTA) model of place field inheritance. A set of place selective cells (“CA3”) project randomly to a set of output neurons (“CA1”). The set of active neurons at the output layer at each position is determined by a KWTA mechanism in which the K most strongly excited neurons are active and all other neurons are silent. In the “no LTP” model, synaptic weights to the output layer slowly randomly fluctuate. In the “LTP” model, weights randomly fluctuate but are also updated according to a Hebbian rule implemented after the KWTA threshold. See “K-Winners-Take-All” section in methods for more details. **B)** LTP/Hebbian plasticity is not necessary for the model to produce place selective cells. Example place selective cells from the output layer of the “LTP” model as in Fig 3E (top) and the “no LTP” model as in Fig 3G (bottom). Each row is a single training iteration of the model, akin to a trial in the real experiment. Columns indicate virtual position. Heatmap indicates the activity rate of the model unit, normalized by the cell’s mean activity across all trials. **C)** The model predicts that cells should have wider place fields and fewer place fields per cell when LTP is intact. *Top-* histogram of single model cell place field width (full width half maximum, full width half maximum, FWHM). *Bottom-*number of place fields per cell (black-LTP model, red-no LTP model) **D)** The model predicts that LTP is not required for stable population coding. The activity of all place selective units is plotted for the first (left) and last trial (right). Cells are sorted by their location of peak activity on the last trial (top - LTP model, bottom - no LTP model). Each row indicates the z-scored activity rate of a unit as a function of position as in Fig 3F & H. The intact representation on trial 1 indicates that LTP stabilizes and exaggerates differences in the random connectivity of feedforward projections. **E)** Our *in vivo* data corroborate the findings from the model. Control animals have wider place fields than Cre animals. Normalized histogram of place field widths (FWHM) for day 1 data combined across all mice (control-black, Cre-red, top-familiar, bottom-novel) **F)** *In vivo* CA1 LTP increases place field width. Within animal average field width (FWHM of peak activity for each place cell) on each day. [N=9 control mice, 7 Cre mice, 6 days. Left - familiar trials mixed effects ANOVA: virus main effect p=5.25×10^−5^, day main effect p=0.015, interaction p=0.467. Right - novel trials mixed effects ANOVA: virus main effect p=6.10×10^−4^, day main effect p=3.69×10^−4^, interaction p=0.13] **G)** Control animals have fewer place fields per cell than Cre animals. Normalized histogram of number of place fields per cell for day 1 data combined across all mice (control-black, Cre-red, top-familiar, bottom-novel) **H)** *In vivo* CA1 LTP decreases the number of place fields per cell. Within animal average number of fields per cell on each day. [N=9 control mice, 7 Cre mice. Left - familiar trials mixed effects ANOVA: virus main effect p=1.16×10^−4^, day main effect p=5.92×10^−4^, interaction p=0.276. Right - novel trials mixed effects ANOVA: virus main effect p=1.14×10^−5^, day main effect p=8.71×10^−3^, interaction p=0.299]. While significant, the effects described in panels E-H did not affect the ability to decode position from neural activity (Fig S11), suggesting that there is enough information for a downstream system to read out a spatial map. *** indicates p<.001*** indicate p<.001

**Fig. S15.**
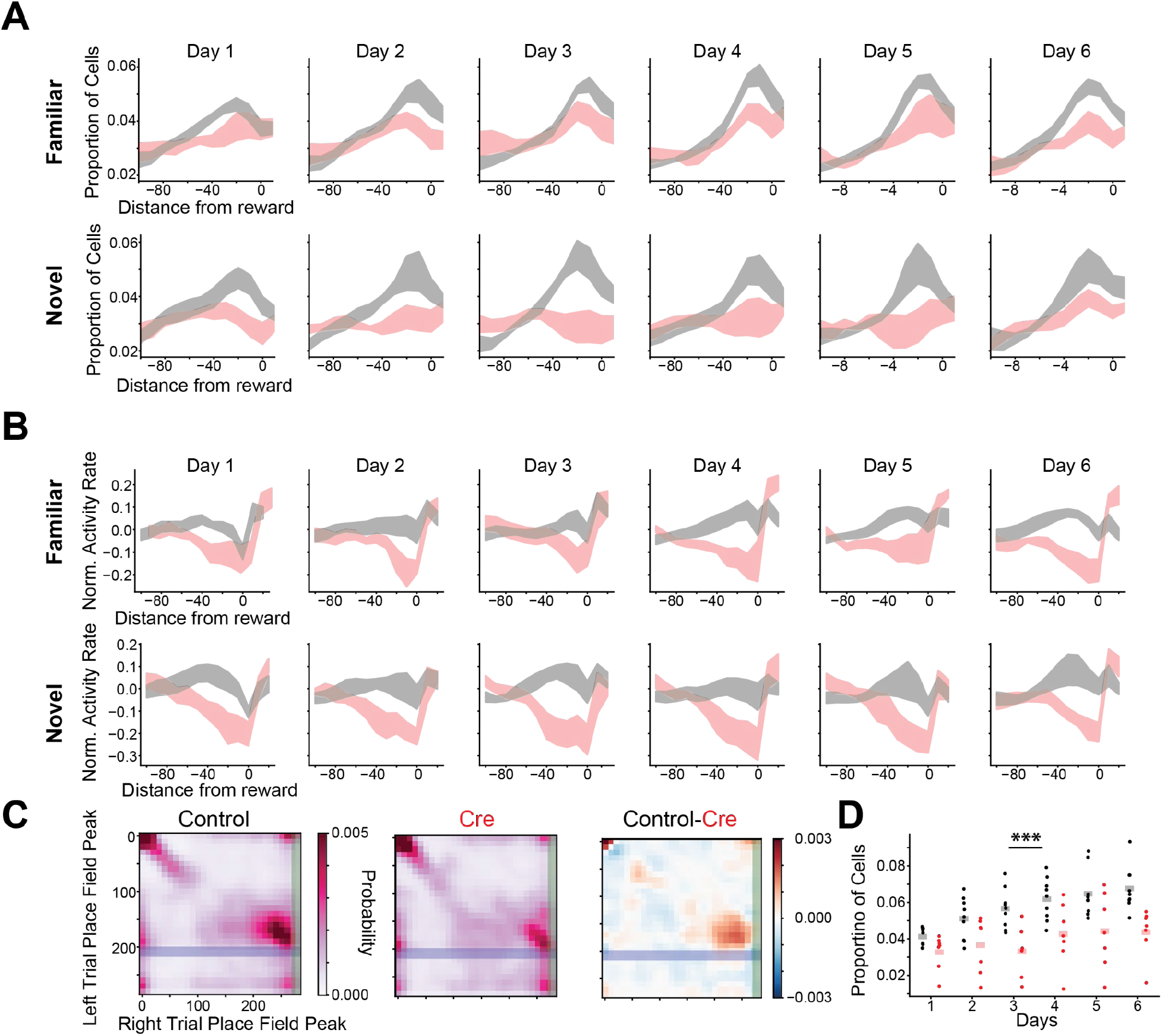
CA1 LTP is required for typical population codes for rewarded locations. **A)** LTP is required for over-representation of reward locations. Proportion of place cells that have their peak in the peri-reward spatial bins for each day (top-familiar, bottom-novel). Data are plotted as across animal mean ± SEM (black-control, red-Cre). **B)** As an alternative to place cell proportion, we quantified reward representation by calculating a spatially normalized population activity rate. We z-scored each cell’s trial-averaged activity rate map as in Fig 3F, 3H & 4A and took the average of that value across all cells, giving one value for each position for each mouse (i.e. average the rows of Fig 4A). We saw a similar effect as the place cell proportion results where the peri-reward activity rate is higher in control animals than Cre animals for both familiar trials [Top - N=9 control mice, 7 Cre mice, 6 days. mixed effects ANOVA: virus main effect p=0.001, day main effect p=0.558, interaction p=0.103] and novel trials [Bottom - N=9 control mice, 7 Cre mice, 6 days. mixed effects ANOVA: virus main effect p=0.001, day main effect p=0.558, interaction p=0.168]. The drop in activity rate in Cre animals correlates with slowing down for rewards (Fig S9). **C)** A greater proportion of cells code for approach to reward locations in control animals than Cre animals. We show a joint normalized histogram of the locations of peak activity on left and right trials for place cells on day 6 (left - control, middle - Cre). Reward locations are highlighted by the shaded regions (blue - left reward zone, green - right reward zone). If a cell does not remap across trials it will lie on the diagonal (i.e. the place field peak is in the same location in both left and right trials). If a cell remaps specifically to match shifts in reward locations (i.e. it codes for approach to rewards in a context invariant manner) it will lie on the off-diagonal near the intersection of reward zones (“reward cells”). The difference between control and Cre histograms (right) highlights more reward cells in the control mice. **D)** Proportion of “reward cells” is shown for each mouse on each day. [N=9 control, 7 Cre mice, 6 days, Mixed effects ANOVA: virus main effect p=6.52×10^−4^, day main effect p=2.37×10^−24^, interaction p=9.89×10^−7^. Posthoc pairwise t-tests (Holm corrected): day 5 control vs Cre p=0.003, day 6 control vs Cre p=0.001, all other days p>.05]

**Fig. S16.**
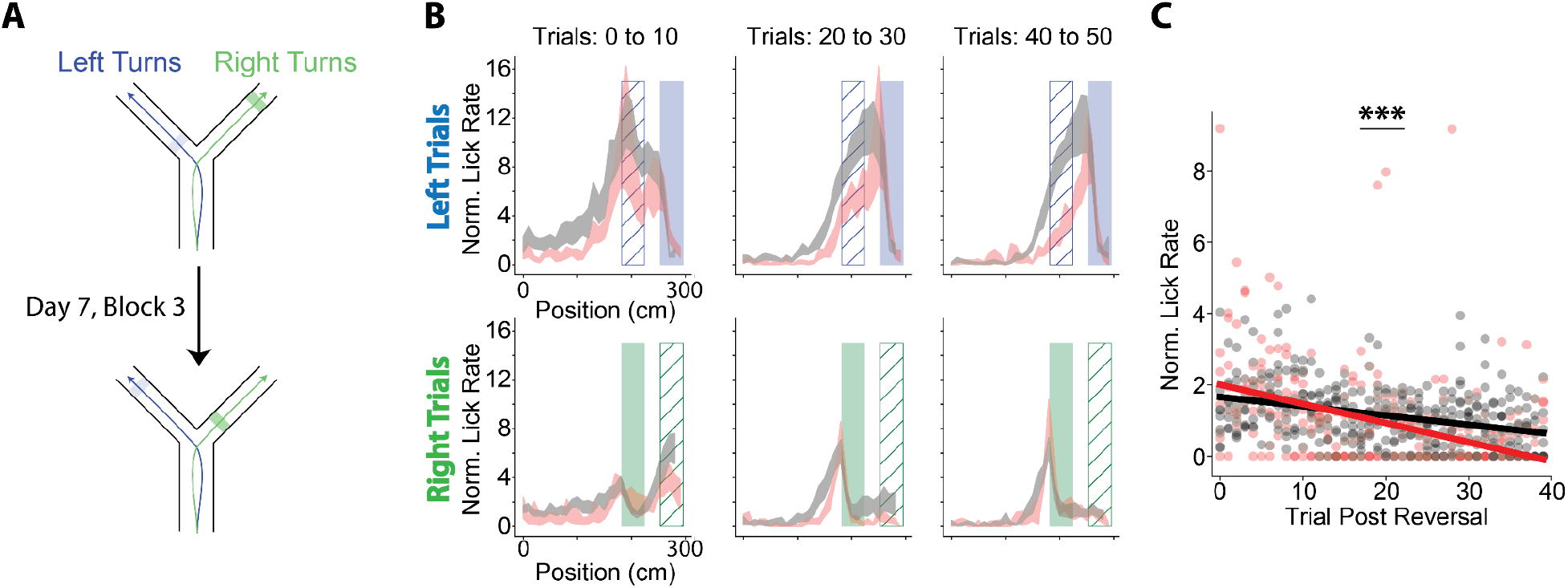
Mice lacking CA1 LTP extinguish licking faster after rewards are moved. **A)** Schematic of the reward reversal task. On day 7, the animals performed the first 2 blocks of trials as in day 6 (randomly interleaved left and right trials). At the beginning of block 3, the reward locations on the two arms were switched. The rewards stayed in this new location for the remaining trials on day 7 and all trials on day 8. **B)** Normalized licking rate as a function of position for binned trials following the reward reversal (top - left trials, bottom -right trials). Each animal’s lick rate is divided by the mean lick rate on the 10 trials preceding the reversal (9 control mice, 6 Cre mice). Data are shown as across animal mean ± SEM. **C)** Normalized lick rate in the previous reward zone is plotted as a function of trials post reversal. Best fit lines from a linear mixed effects model are shown (random intercepts linear mixed effects model: virus main effect p=0.157, trial main effect p=2.36×10^−8^ virus x trial interaction p = 5.22×10^−4^)

**Fig. S17.**
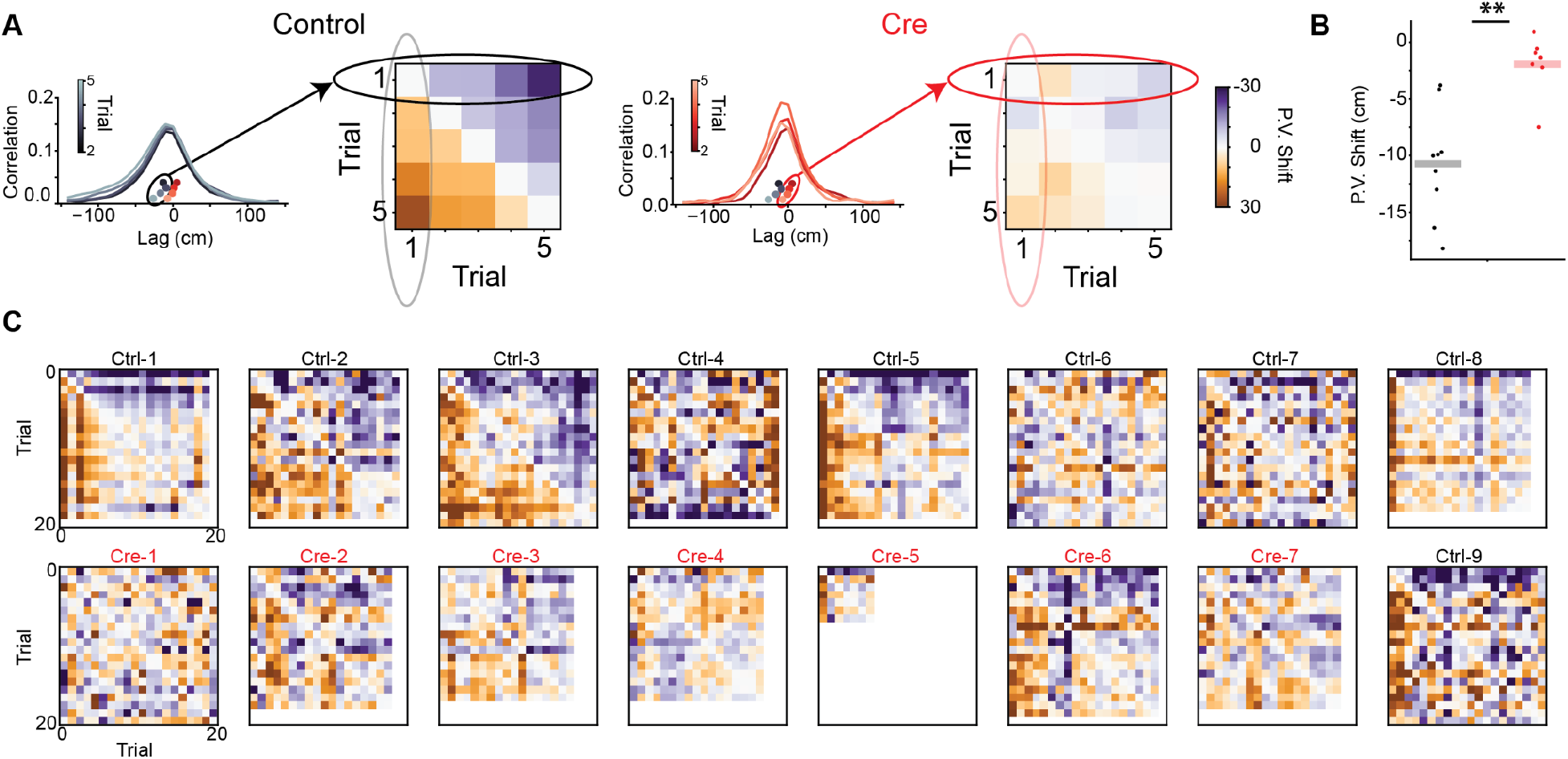
Abolishing LTP reduces the backward shift of population place codes in the initial experience of novel environments. **A)** Schematic showing how the population vector (PV) shift is calculated. For each mouse we calculate the center of mass from trial x trial PV cross correlations. In the cross-correlation plots, reproduced from Fig 4D, the cross correlations of each trial with trial 1 are shown. These center of mass values yield a trial x trial antisymmetric PV shift matrix. The trial 1 examples give the entries in the first row/column of the matrix. The matrices shown are the across animal average PV shifts for each pair of trials. The average PV shift is calculated by averaging the upper triangle of this matrix. All data are from day 1 novel trials. **B)** Control mice display a greater backward PV shift than Cre mice. Replotted day 1 PV shift from Fig 4E. Posthoc t-test p=2.44×10^−3^ (Holm corrected) **C)** PV shift matrices for all novel arm trials on day 1 for all mice. White space indicates trials that were not completed due to slower running speeds or stopping. Note that PV shift statistics are calculated using only the first 5 trials.

**Fig. S18.**
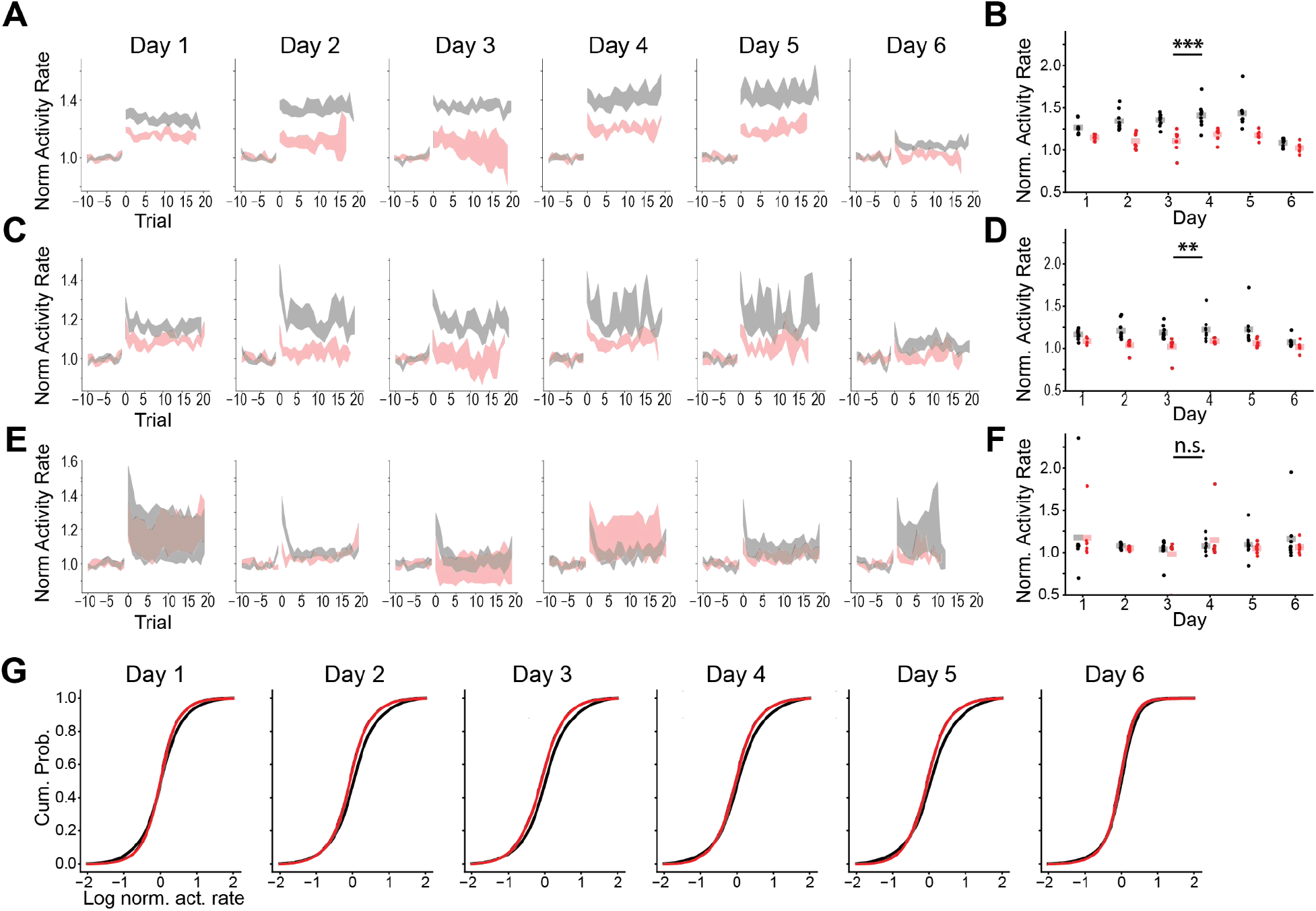
LTP is necessary for increased activity rates in novel environments. **A)** Population averaged normalized activity rate for novel trials in the last trial block on each day. Shaded region indicates mouse average ± SEM. Note: day 6 has equivalent “familiar” and “novel” trials, acting as an internal control for novelty. **B)** Averaged activity rate on novel arm trials for each mouse on each day. Shaded bars indicate across animal means. [N = 9 control, 7 Cre mice, 5 days. Mixed effects ANOVA: virus main effect p=4.59×10^−5^, day main effect p=0.005, interaction p=0.208] **C)** Same as (A) for familiar trials in the last trial block on each day. **D)** Same as (B) for trials shown in (C). Control animals also show increased activity rate in familiar trials in the last block compared to Cre animals. Given that novel and familiar trials are interleaved during this block, this indicates that the novel arm exposure alone (or the removal of the translucent curtain during this block) is sufficient to generalize the activity rate increase. However, the magnitude of the effect is smaller than the response to novel trials. [N=9 control, 7 Cre mice, 5 days. Mixed effects ANOVA: virus main effect p=0.002, day main effect p=0.498, interaction p=0.513] **E)** Same as (A) for the block 3 to block 4 transition on each day. This analysis is shown as an internal control as there is no novel experience in this transition. **F)** Same as (B) for trials shown in (E). This strengthens our conclusion in (D) that the increase seen in control mice is a carry-over from the interleaved novel trials (or removal of the virtual curtain). [N=9 control, 7 Cre mice, 5 days. Mixed effects ANOVA: virus main effect p=0.830, day main effect p=0.22, interaction p=0.904] **G)** Cumulative probability of each cell’s log-transformed normalized activity rate in novel trials on each day. To weigh mice evenly, cumulative probabilities are calculated for each mouse independently and then averaged across mice within a group (black-control, red-Cre) before being renormalized to be a valid cumulative mass function. Note that the increase in activity is shared across a large proportion of the population.

